# Arousal state modulates human hippocampal ripples

**DOI:** 10.64898/2026.06.26.734578

**Authors:** Elizabeth M. Siefert, Yvonne Y. Chen, Kathryn A. Davis, Han-Chiao I. Chen, Anna C. Schapiro, Brett L. Foster

## Abstract

Hippocampal ripples are transient, high-frequency oscillations linked to memory replay and consolidation. Ripples are well-characterized in rodents to occur during periods of behavioral inactivity (i.e., sleep, rest), viewed as “offline” states where replay can emerge with limited sensory interference. However, human studies have increasingly observed ripples during active tasks, raising the questions of whether ripple genesis and function have been misunderstood or whether there are fundamental species differences. We propose that low arousal states—predominant during offline sleep and transient during wake—may constitute a common mechanism of ripple genesis that reconciles these observations. We recorded directly from human hippocampus during sleep and wake, measuring arousal via sleep staging, pupillometry, and heart rate. Ripple occurrence consistently tracked low arousal: rates were maximal in NREM sleep, small-pupil wake states, and slow heart rate periods across sleep and wake. This modulation was stronger in anterior than posterior hippocampus and was hippocampus specific: ripple-like activity outside the hippocampus showed an opposite modulation, increasing with high arousal. These results resolve apparent species differences and provide a unifying view of offline periods as arousal dips that can emerge across behavioral states, including transiently during active wake, suggesting hippocampal ripples, and memory consolidation, occur continuously intermixed with cognition.

## Introduction

Memory formation is a process that extends far beyond the initial moment of encoding. After experiences are first encoded in the hippocampus, their transformation from transient representations into enduring memories unfolds over timescales ranging from minutes to years. A key mechanism underlying this consolidation process is memory replay, the repeated reactivation of hippocampal memory representations, thought to drive coordinated cortical reinstatement and, over time, the formation of distributed cortical memory representations (Born & Wilhelm, 2012; McClelland et al., 1995). In rodents, replay has predominantly been observed during sleep and awake rest (Kudrimoti et al., 1999; A. K. Lee & Wilson, 2002; Skaggs & McNaughton, 1996; Wilson & McNaughton, 1994), as these “offline” periods of inactivity are thought to provide privileged moments for internal replay to transform memory representations while limiting external interference from new sensory information (Hasselmo, 2006; Klinzing et al., 2019; Mednick et al., 2011).

Hippocampal memory replay can be studied through the detection of ripples—transient, high-frequency oscillations in the local field potential (LFP) that correspond to the local circuit firing during memory replay spike sequences (Buzsáki, 2015; Girardeau & Zugaro, 2011). In rodents, disruptions to ripples and ripple-associated activity during periods of sleep or inactivity impair memory (Aleman-Zapata et al., 2022; Ego-Stengel & Wilson, 2010; Girardeau et al., 2009; Y. Zhang et al., 2021). Consistent with rodent findings, human studies have shown that hippocampal ripples occur prominently during sleep, coordinated with other memory-related sleep oscillations (Ngo et al., 2020; Schreiner et al., 2024; Skelin et al., 2021; Staresina et al., 2015, 2023). These observations support the use of ripples as a cross-species marker of replay and underscore the importance of offline states for memory consolidation. However, a recent body of work has suggested human hippocampal ripples occur during active wakefulness, as ripples have been observed during cognitive tasks ranging from perception (Y. Y. Chen et al., 2021), to memory (both encoding and retrieval; Henin et al., 2021; Norman et al., 2019; Sakon et al., 2024; Sakon & Kahana, 2022; Vaz et al., 2019) to planning and decision making (He et al., 2026; Zhou et al., 2026), with rates in some cases exceeding those typically reported during sleep. The observance of ripples during active tasks in humans stands in tension with the rodent literature, where they are predominantly associated with rest and inactivity, complicating the tight coupling of ripples—and by proxy replay and consolidation—with offline states. Further challenging the cross-species comparison, while rodent studies have focused largely on hippocampal ripples specifically, in humans ripples have also been studied in the amygdala and broader medial temporal lobe (MTL) structures (Clemens et al., 2007; Cox et al., 2020; Sakon et al., 2024; Sakon & Kahana, 2022; Staresina et al., 2023; Vaz et al., 2019; H. Zhang et al., 2024), as well as in neocortical regions (Dickey et al., 2022; Verzhbinsky et al., 2024). Thus, the human ripple literature appears to diverge from the rodent in both the behavioral context (offline sleep/rest vs. online wake) and anatomical location (hippocampus vs. broader MTL) in which ripples are reported. These observations raise a fundamental question: what determines when hippocampal ripples occur? Is there a unifying mechanism that could bring together the rodent and human literature—one that accounts for their high rates during offline states like sleep while also explaining their occurrence during active waking tasks?

Given the tight coupling of hippocampal ripple events with offline sleep states in rodents, one candidate mechanism governing their occurrence is arousal. Consistent with this view, ripples do not occur uniformly across sleep, but instead co-vary with arousal level across sleep stages, with rates greatly elevated in NREM and attenuated in REM (Buzsáki, 2015; Joo & Frank, 2018). Furthermore, within NREM sleep, ripple occurrence has been tightly linked to reductions in arousal in rodents, as indexed by momentary decreases in acetylcholine (Y. Zhang et al., 2021, 2024) and pupil diameter (H. Chang et al., 2025). This relationship between hippocampal ripple events and pupil-linked arousal was also observed previously during waking in rodents (Farrell et al., 2024; McGinley, David, et al., 2015; Nguyen et al., 2024). Together, these findings from rodents suggest that low arousal promotes ripple genesis. Yet, whether this arousal-ripple relationship holds in humans remains an open question. Critically, while arousal drops substantially during sleep, it also fluctuates dynamically during wakefulness and while performing cognitive tasks. This raises the possibility that, if arousal similarly governs ripple occurrence in humans, it may serve as a unifying variable across species and behavioral states: sustained low arousal during sleep could account for the predominance of ripples during sleep in rodents, while transient arousal dips during active tasks could drive their occurrence during wakefulness in humans. Under this framework, offline states would be reframed not as overt behavioral inactivity, but instead as periods of low arousal that can occur across both sleep and wake, with ripple rates increasing with the degree of arousal reduction more generally.

Here, we tested the relationship between arousal and hippocampal ripple genesis in humans. Using invasive intracranial recordings in humans, we measured hippocampal ripples during both overnight sleep and separate periods of wake, indexing arousal via sleep staging and pupillometry, respectively. Across sleep and wake recordings, a consistent result emerged: hippocampal ripples occurred preferentially during states of low arousal, with elevated ripple rates during NREM in sleep and small pupil states in wake. To bridge across recordings, we leveraged heart rate—a measure of arousal obtainable during both sleep and wake—and replicated the finding, demonstrating that ripples occur preferentially during periods of low heart rate regardless of behavioral state. Finally, we showed that this relationship is anatomically specific to the hippocampus and strongest anteriorly: Ripple events in the anterior hippocampus consistently demonstrated tighter coupling to arousal than the posterior hippocampus, while ripple-like events in the amygdala showed the opposite pattern, with rates increasing during periods of high arousal. Together, these results offer a parsimonious account of when and why hippocampal ripples occur in humans, one that reconciles the human and rodent literatures, reframes offline states as dynamic periods of low arousal, and connects ripple occurrence to underlying mechanisms of arousal physiology.

## Results

We hypothesized that hippocampal ripple rates would increase during low arousal states, tracking both large arousal drops across sleep stages and subtler dips within wakefulness (Fig. 1a). To test this, we performed human intracranial recordings in 20 neurosurgical patients during both overnight sleep and a wakeful fixation task with pupillometry (Fig. 1; see Methods). Recordings targeted the hippocampus (119 electrodes) and amygdala (77 amygdala electrodes; Fig. 1b; Table S1). The hippocampus was the main site of investigation, as it is the primary locus of ripple genesis, and was subdivided into anterior (69 electrodes) and posterior (50 electrodes) regions, as these areas are known to differ functionally (Moser & Moser, 1998; Poppenk et al., 2013; Strange et al., 2014; Fig. 1b). Amygdala recordings were analyzed given recent human studies reporting amygdala ripple activity (Cox et al., 2020; Sakon et al., 2024; Staresina et al., 2023; H. Zhang et al., 2024), and to provide a proximal MTL comparison region for hippocampal effects. Ripple events were detected across anatomical regions using an established detection and spectral verification method (Y. Y. Chen et al., 2021), following consensus guidelines (A. A. Liu et al., 2022; Fig. 1c). This approach detects putative ripple events and then confirms spectral evidence of ripple band oscillations, excluding other broadband events which could also drive increases in ripple band power (e.g., broadband gamma, interictal spikes). Ripple events were examined across levels of arousal during sleep via staging (e.g., NREM vs. wake) and during wake via pupillometry (e.g., diameter small vs. large; Fig. 1d).

**Figure 1.**
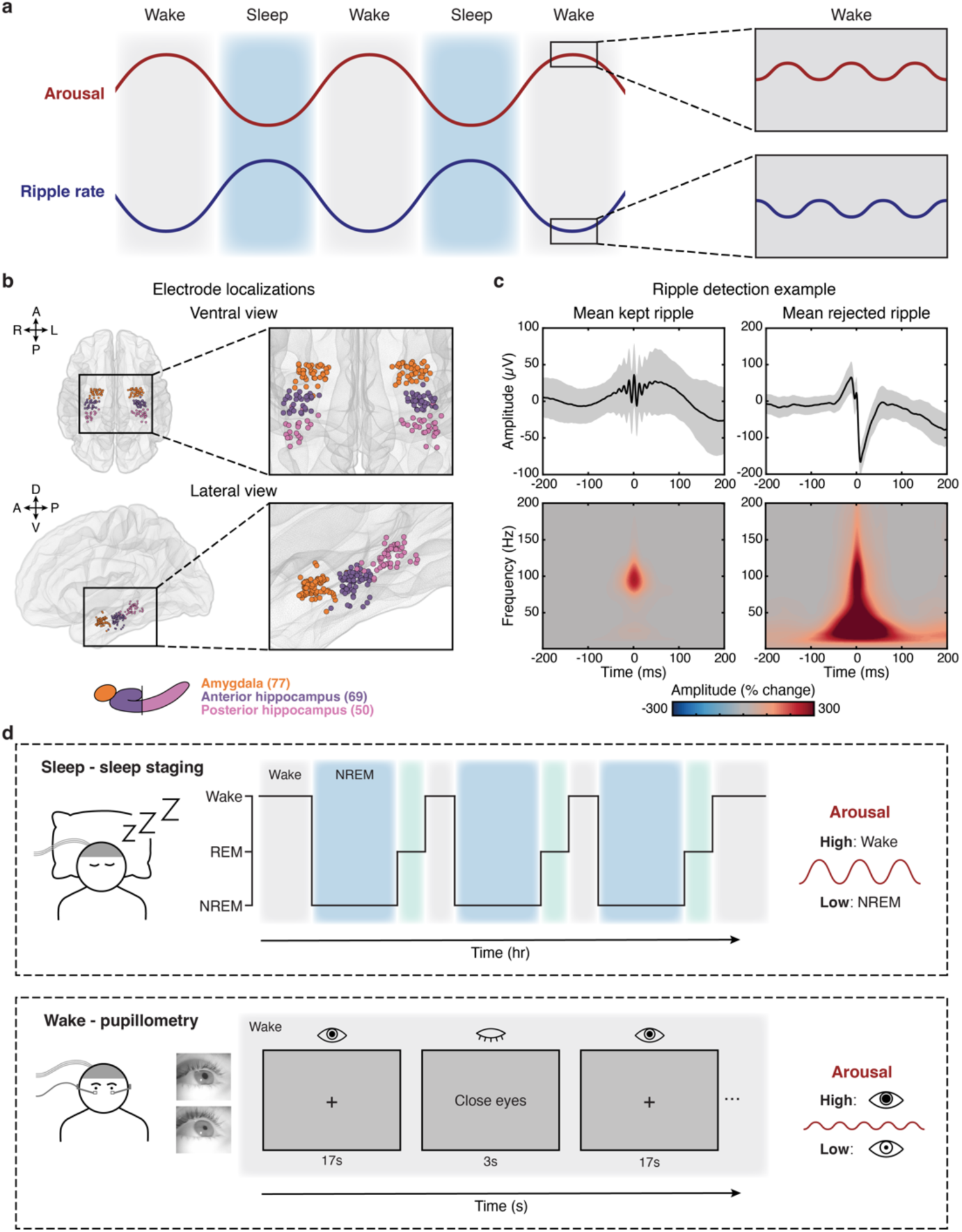
Experimental overview. **a.** Framework: large arousal fluctuations (red) across sleep and wake are accompanied by corresponding fluctuations in hippocampal ripple rate (blue), with ripples most frequent during low arousal states like sleep. Within each state, more subtle arousal fluctuations similarly covary with ripple rate, even during waking when overall ripple rates are reduced. **b.** Intracranial electrode locations for all participants (n = 20), normalized to MNI space, viewed from the ventral surface (top) and lateral surface (hemispheres collapsed; bottom). Colors indicate brain region: amygdala (orange), anterior hippocampus (ant. hipp., purple), posterior hippocampus (post. hipp., pink); electrode counts in parentheses. **c.** Mean LFP voltage trace (top; shading ± SD) and time-frequency representation (bottom) for kept (left) and rejected (right) hippocampal ripple events from a representative sleep session (P4). **d.** Multiple nights of sleep and blocks of a waking fixation task were collected from all participants. Sleep (top): arousal was indexed via sleep staging (Wake: high arousal; NREM: low arousal). Wake fixation task (bottom): arousal was measured via pupillometry using a custom head-mounted system. Example eye camera frames show moments of large and small pupil size. During the task, participants fixated for ∼17s, closed their eyes for ∼3s, and reopened them on a tone cue (10 or 15 trials per block). Pupil size during eyes-open periods indexed arousal (larger pupil: higher arousal; smaller: lower).

### Hippocampal ripple rate increased during NREM states in sleep

We first assessed the relationship between hippocampal ripple events and arousal fluctuations during overnight sleep. Arousal was measured by sleep staging, with NREM and wake reflecting low and high arousal states, respectively. Two nights of sleep were recorded and scored from each participant, except for one participant for whom only one night was included (41 total nights; see Methods; Table S1). Participants slept on average 7.5 h (SD = 1.7 h), with each night containing at least 1 hr of NREM and 1 hr of wake (Fig. S1; Table S2, S3). To test the influence of sleep stage on hippocampal ripple rate, we employed a linear mixed-effects model with fixed effects of sleep stage (NREM, REM, wake) and region (anterior, posterior) and random intercepts for electrodes nested within participants. Across the hippocampus, sleep stage predicted ripple rate (main effect of sleep stage: χ²(2) = 405.78, *p* < 0.001; Fig. 2d), with ripple rates highest during NREM (model mean ± SEM = 0.050 ± 0.002 Hz), reduced during REM (0.035 ± 0.002 Hz), and lowest during wake (0.032 ± 0.002 Hz; NREM – REM: *t*(577) = 11.66, *p* < 0.001; NREM – wake: *t*(575) = 14.76, *p* < 0.001; REM – wake: *t*(577) = 2.89, *p* = 0.011; all Tukey-corrected; Fig. 2d), consistent with prior reports in rodents and humans (Buzsáki, 2015; Y. Y. Chen et al., 2021; Jiang et al., 2020; Joo & Frank, 2018).

**Figure 2.**
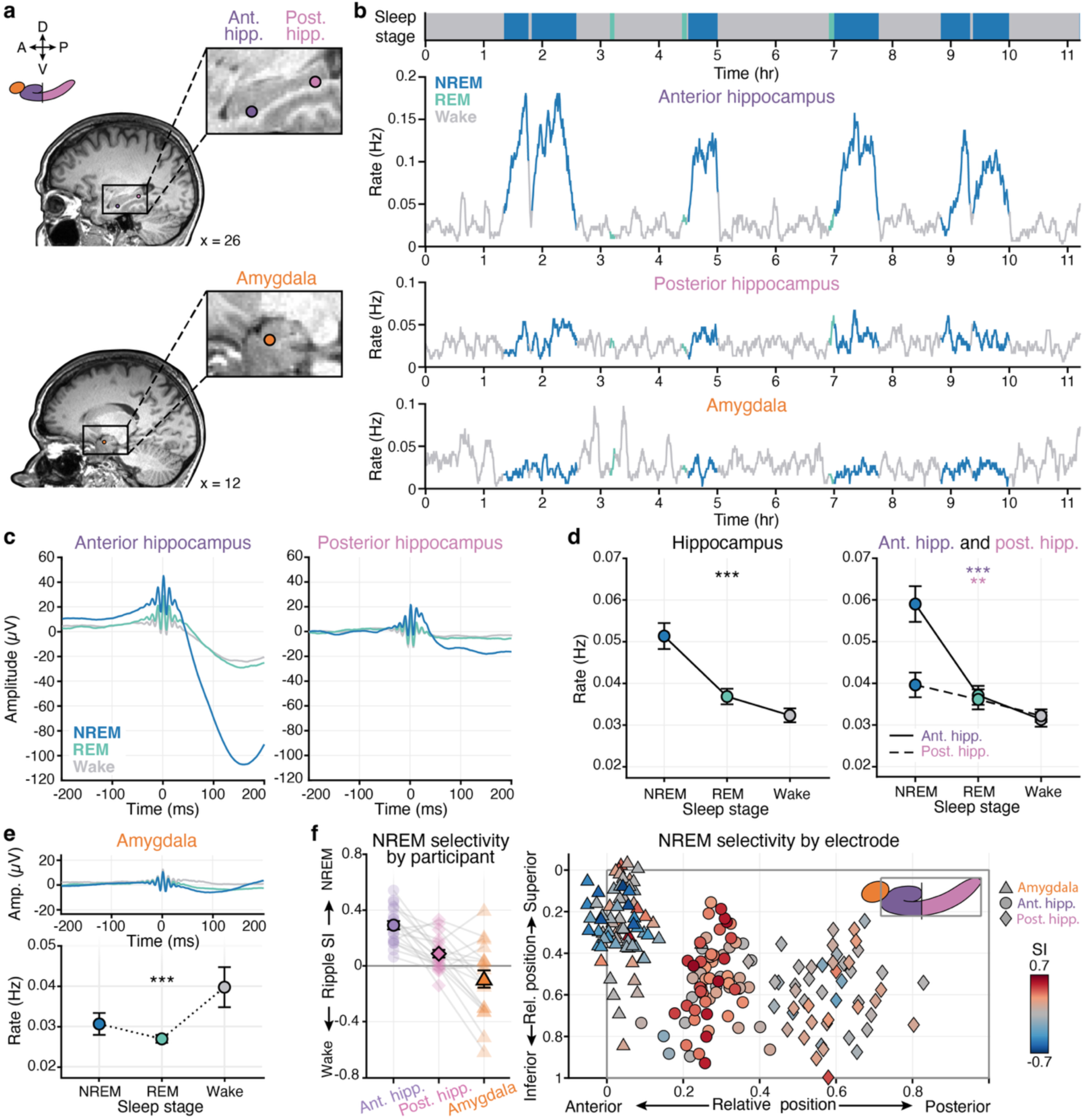
Ripple rate across sleep stages. **a.** Example electrodes located in anterior hippocampus (purple), posterior hippocampus (pink), and amygdala (orange) from a single participant (all right hemisphere; P11). **b.** Ripple rate across a representative sleep night for the electrodes in a. Top bar: sleep stages over time (NREM: dark blue; REM: green; Wake: grey). **c.** Group mean ripple-triggered voltage traces per sleep stage for anterior hippocampus (left) and posterior hippocampus (right). **d.** Group mean ripple rate per sleep stage for all hippocampal electrodes (right), and separately for anterior hippocampus (solid) and posterior hippocampus (dashed) (left). **e.** Group mean ripple-triggered voltage traces (top) and ripple rate (bottom) per sleep stage for amygdala electrodes. **f.** Ripple rate selectivity index (SI) to sleep stage (positive values = more ripples during NREM than Wake = selectivity to NREM; negative values = selectivity to Wake). Left: SI by anatomical region for individual participants (transparent shapes) and group mean (solid shapes). Right: Individual electrodes plotted by anatomical position relative to the most anterior and superior point of each participant’s hippocampus, colored by SI (red: selectivity to NREM; blue: selectivity to Wake); shapes indicate anatomical region (anterior hippocampus: circle; posterior hippocampus: diamond; amygdala: triangle). For **d, e** filled circles: group mean; error bars: ±SEM across group; significance of main effect of sleep stage: \*\*\**p* < 0.001, \*\**p* < 0.01.

While increased hippocampal ripple rates during NREM has previously been observed across species, recent evidence suggests there may be important differences in hippocampal ripple activity across the long axis in humans (De Filippo & Schmitz, 2023; Jiang et al., 2020; Staresina et al., 2023; To et al., 2025; J. Zhang et al., 2025). We thus sought to examine if there were any differences in arousal-ripple modulation between anterior and posterior hippocampus. To test this, we evaluated the effect of sleep stage on ripple rate separately for each hippocampal region (anterior, posterior), as well as the interaction between region and sleep stage, in the mixed-effects model described above. Ripple rate tracked sleep stage in both anterior and posterior hippocampus (joint F-tests: anterior: F(2, 575.37) = 201.47, *p* < 0.001; posterior: F(2, 575.37) = 6.27, *p* = 0.002), with rates higher in NREM than wake in both regions (Fig. 2b, d; anterior hippocampus: NREM–wake: t(572) = 18.79, *p* < 0.001; NREM–REM: t(574) = 15.62, *p* < 0.001; REM–wake: t(574) = 2.99, *p* = 0.008; posterior hippocampus: NREM–wake: t(572) = 3.51, *p* = 0.001; NREM–REM: t(575) = 2.18, *p* = 0.075; REM–wake: t(575) = 1.27, *p* = 0.412; all Tukey-corrected). However, the magnitude of this modulation was significantly greater—3x larger—in anterior than posterior hippocampus (sleep stage × region: χ²(2) = 107.22, *p* < 0.001), driven by substantially elevated ripple rates in anterior than posterior hippocampus during NREM (*t*(205) = 8.88, *p* < 0.001; Tukey-corrected; Fig. 2b, d). Rates in the two regions were matched during REM (*t*(213) = 1.12, *p* = 0.266) and wake (*t*(205) = 0.25, *p* = 0.801), suggesting the anterior-posterior NREM difference reflected a functional or mechanistic distinction rather than a measurement difference. Additionally, ripple events in anterior hippocampus showed a pronounced post-ripple wave (PRW, Hussin et al., 2020) in the LFP that appeared largest in NREM (Fig. 2c). To quantify this effect, we computed the mean LFP amplitude in a 50 to 150ms post-ripple window baselined to a -150 to -50ms pre-ripple window and evaluated how it was impacted by sleep stage (NREM, REM, wake) and region (anterior, posterior) with a mixed-effects model. Anterior hippocampal PRW-amplitude varied across sleep stages and was greatest in magnitude in NREM (F(2, 237.09) = 20.91, *p* < 0.001; NREM – REM: t(233) = –6.46, *p* < 0.001; NREM – wake: t(233) = –4.00, *p* < 0.001; REM – wake: t(233) = 2.50, *p* = 0.035; all Tukey-corrected; Fig. 2c). This effect in anterior hippocampus was stronger than that in posterior hippocampus (region: χ²(1) = 18.26, *p* < 0.001; region × sleep stage: χ²(2) = 15.30, *p* < 0.001), where the PRW did not significantly vary across sleep stages (F(2, 237.09) = 0.31, *p* = 0.733; Fig. 2c).

The amygdala showed a strikingly different pattern: amygdala ripple-like event rates were highest during wake (mixed-effects model with region as hippocampus or amygdala; amygdala: F(2, 946.83) = 19.17, *p* < 0.001; wake: 0.038 ± 0.002 Hz, NREM: 0.033 ± 0.002 Hz, REM: 0.028 ± 0.002 Hz; NREM – REM: t(942) = 2.89, *p* = 0.011; NREM – wake: t(938) = −3.34, *p* = 0.003; REM – wake: t(942) = −6.20, *p* < 0.001; all Tukey-corrected). This sleep stage modulation differed significantly from that of the entire hippocampus (sleep stage × region: χ²(2) = 156.75, *p* < 0.001), with overall event rates also lower in the amygdala (region: χ²(1) = 86.60, *p* < 0.001). Ripple-like events in the amygdala had a significantly smaller PRW (region: χ²(1) = 7.41, *p* = 0.006) and a smaller sleep stage-PRW effect than the entire hippocampus (region × stage: χ²(2) = 7.01, *p* = 0.030), as there was no difference in the amygdala PRW across sleep stages (amygdala: F(2, 388.1) = 1.40, *p* = 0.248; Fig. 2e). Given the diverging results between hippocampus and amygdala, we next asked whether the amygdala effect was structure-specific or reflected broader non-hippocampal activity. To address this, we detected ripple-like events in a set of lateral temporal electrodes from the same probes used for hippocampal and amygdala analyses and examined their sleep stage modulation (n = 20, 78 lateral temporal electrodes; see Methods; Fig. S5). Ripple-like events detected at cortical sites displayed a sleep stage modulation profile similar to that of the amygdala, with rates highest in wake, and did not differ significantly from the amygdala at any stage (F(2, 1322.83) = 15.82, p < 0.001; NREM – REM: t(1320) = 1.52, *p* = 0.284; NREM – wake: t(1314) = –3.98, *p* < 0.001; REM – wake: t(1320) = –5.44, *p* < 0.001; amygdala – cortex, all t < 0.44, *p* > 0.900; Tukey-corrected; Fig. S5), suggesting the amygdala pattern was not structure specific. This observation supports the use of the amygdala as a control region and further highlights the anatomical specificity of hippocampal ripple-sleep stage modulation. All of the described trends replicated when sleep stages were resolved on a more granular level (N1, N2, N3, REM, wake): anterior hippocampal ripple rates were highest in N3, decreased progressively from N3 to N2 to N1, and were lowest during REM and wake, with the posterior hippocampus demonstrating an attenuated version of this pattern, whereas the amygdala showed no comparable modulation (Fig. S2).

Motivated by the striking increase in hippocampal ripple rate during NREM sleep, which was more pronounced in anterior than posterior hippocampus, we next investigated how this NREM-ripple modulation varied continuously along the hippocampal long axis. For each electrode, we quantified the degree to which ripple rates increased during NREM compared to wake by computing a NREM selectivity index (SI = (NREM rate – wake rate) / (NREM rate + wake rate), where positive and negative values reflected greater ripple rates during NREM and wake, respectively. Electrodes were colored by their SI value and plotted in a normalized hippocampal space across participants, along the anterior-posterior (AP) and inferior-superior (IS) anatomical axes (Fig. 2f; see Methods). NREM selectivity was strongest in the most anterior hippocampal electrodes with SI values decreasing toward posterior hippocampus (Fig 2f; mixed effects model predicting SI as a function of AP position: slope (anterior to posterior) = –0.641 ± 0.085, t(109.84) = −7.57, *p* < 0.001). Replicating this effect, in a mixed-effects model predicting ripple rate as a function of sleep stage and continuous AP position, a significant sleep stage × AP position interaction emerged (χ²(2) = 87.14, *p* < 0.001), with NREM ripple rates highest in anterior hippocampus and decreasing posteriorly (slope (anterior to posterior) = –0.067 ± 0.008 Hz, t(209) = −8.46, *p* < 0.001). Together, these results demonstrate that hippocampal ripple genesis is favored during the low-arousal state of NREM, an effect specific to the hippocampus and strongest anteriorly.

### Hippocampal ripple rate increased during small pupil states in wake

Having shown that ripple rates increase to their highest levels during low arousal states in sleep (i.e., NREM), we asked whether a similar relationship holds for low arousal states during wakefulness. In contrast to the large arousal changes that occur across sleep stages, arousal fluctuations within wake are more subtle and transient in nature (McCormick et al., 2020; McGinley, Vinck, et al., 2015). To capture these dynamics, we performed joint sEEG-pupillometry recordings during periods of wake, where small and large pupil diameter indexed low and high arousal, respectively (McGinley, David, et al., 2015; Murphy et al., 2014; Reimer et al., 2014, 2016). However, for simultaneous sEEG-pupillometry recordings, standard desktop methods pose considerable practical challenges, as neurosurgical patients are often reclining in bed and wearing large head wraps. We addressed these constraints with a custom head-mounted system using 3D-printed flexible frames to secure cameras monitoring both eyes, enabling high-fidelity continuous pupil diameter measurement synchronized to neural signals that is tolerant to head position and motion (Fig. 1d, 3a; see Methods). Joint recordings were performed during a controlled wake fixation task where participants alternated between eyes-open fixation periods (∼15-17s) and brief eyes-closed (∼3s) recovery periods (median task blocks per participant: 4, IQR: 4, 6; mean total eyes-open fixation time per participant ± SD: 15.68 min ± 7.36 min; Fig. 1d; Table S1; see Methods). A simple fixation task was chosen to minimize visual transients and thereby isolate endogenous, pupil-linked arousal fluctuations separate from stimulus-driven changes or eye-motion-related artifacts. Finally, experiments were performed in a controlled low-light setting (see Methods).

**Figure 3.**
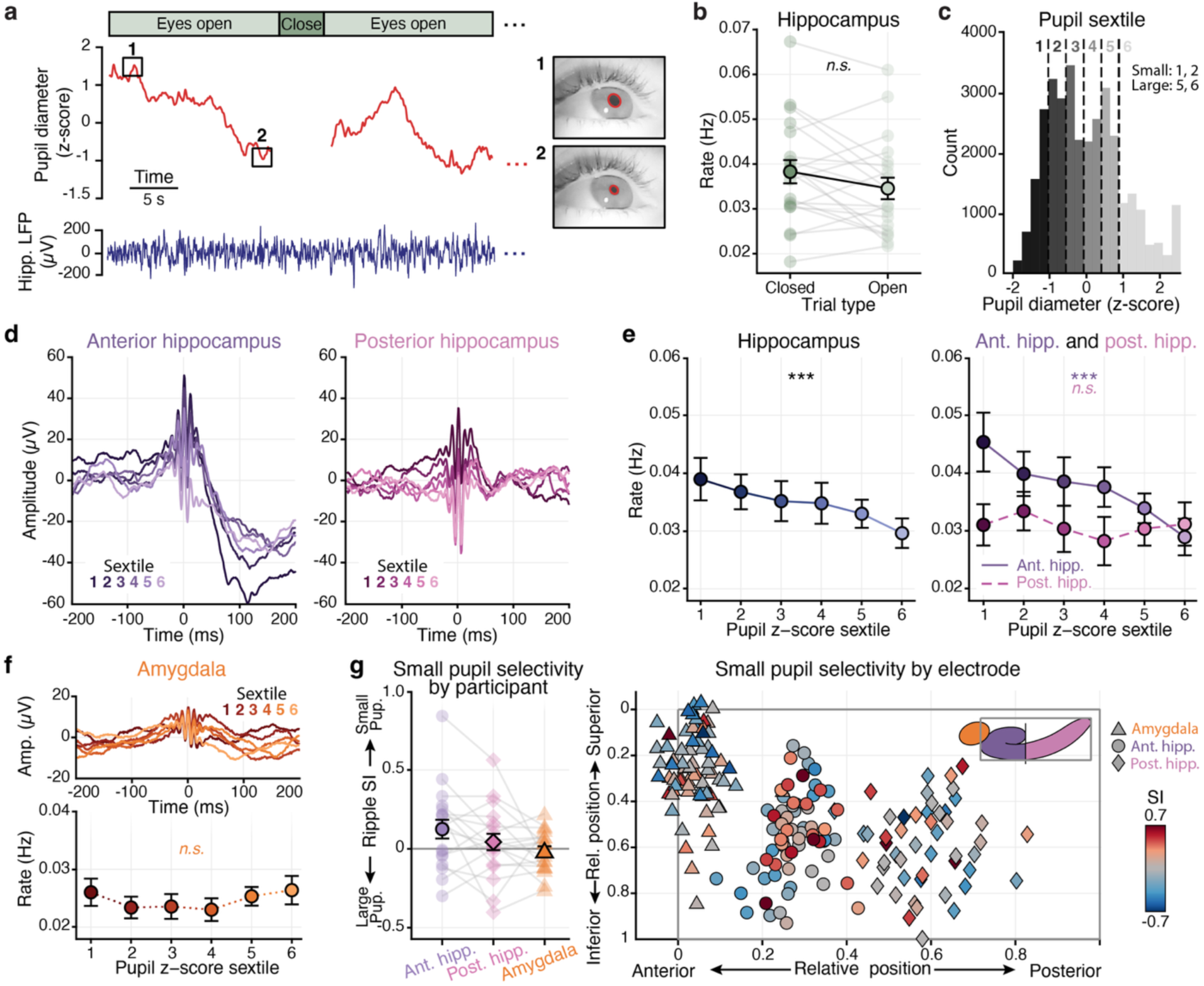
Ripple rate across pupil states. **a.** Example joint hippocampal-pupil recording across two eyes-open fixation phases, with eye video frames from two selected moments (pupil outlined in red; P6). **b.** Mean ripple rate during eyes-closed and eyes-open fixation task periods. **c.** Distribution of pupil diameter from a single task block (P6), divided into sextiles (dark to light: small to large pupil). **d.** Group mean ripple-triggered voltage traces per pupil sextile during the fixation task, for anterior hippocampus (left) and posterior hippocampus (right). Line color indicates sextile (dark to light: sextile 1-6, small to large pupil). **e.** Group mean ripple rate across pupil sextiles for all hippocampal electrodes (right), and separately for anterior hippocampus (solid) and posterior hippocampus (dashed) (left). **f.** Group mean ripple-triggered voltage traces (top) and ripple rate (bottom) across pupil sextiles for amygdala electrodes. **g.** Ripple rate selectivity index (SI) to pupil sextile (positive values = more ripples during small pupil than large pupil states = selectivity to small pupil; negative values = selectivity to large pupil). Left: SI by anatomical region for individual participants (transparent shapes) and group mean (solid shapes). Right: individual electrodes plotted by anatomical position relative to the most anterior and superior point of each participant’s hippocampus, colored by SI (red: selectivity to small pupil; blue: selectivity to large pupil); shapes indicate anatomical region (anterior hippocampus: circle; posterior hippocampus: diamond; amygdala: triangle). For **b, e, f** filled circles: group mean; error bars ±SEM across group; transparent dots and lines: individual participants; significance of main effect of pupil sextile: \*\*\**p* < 0.001.

We examined how arousal fluctuations during wake, as captured by pupil diameter during the eyes-open fixation periods, influenced ripple rates. Overall, in the wake fixation task, mean ripple rate was significantly lower than that observed in NREM sleep (wake fixation task mean rate: 0.035 ± 0.0025 Hz; NREM – wake fixation task: t(412) = 5.18, *p* < 0.001), consistent with a wide literature (Buzsáki, 2015; Y. Y. Chen et al., 2021; Jiang et al., 2020; Joo & Frank, 2018). While mean ripple rate was slightly elevated for eyes-closed periods (Y. Y. Chen et al., 2021), there was no significant difference compared to eyes-open periods (χ²(1) = 3.39, *p* = 0.065; Fig. 3b). In the eyes-open periods, ripple rates were calculated per pupil sextile where sextile 1 corresponded to the smallest pupil diameter (lowest arousal) and sextile 6 to the largest (highest arousal; Fig. 3c), following recent work that linked replay rates to pupil sextile during sleep in the rodent (H. Chang et al., 2025). To evaluate the relationship between pupil sextile and hippocampal ripple events, mixed effects models were applied with predictors of pupil sextile (1-6) and hippocampal region (anterior, posterior). Mirroring our sleep results, during the wake fixation task hippocampal ripple rate was inversely related to arousal state: ripple rates were highest during small pupil (low arousal) states and lowest during large pupil (high arousal) states (main effect of pupil sextile: slope (small to large pupil) = −0.0020 ± 0.0005 Hz, t(593) = −3.91, *p* < 0.001; sextile 1 mean ± SEM: 0.037 ± 0.003 Hz, sextile 6: 0.031 ± 0.003 Hz; Fig. 3e), replicating findings in the rodent (H. Chang et al., 2025; Farrell et al., 2024; McGinley, David, et al., 2015; Nguyen et al., 2024). As with the sleep-stage analyses, we then asked whether this effect differed between anterior and posterior hippocampus and was specific to the hippocampus relative to the amygdala.

Ripple rate inversely tracked pupil sextile in anterior hippocampus (slope = −0.0020 ± 0.0005 Hz, t(593) = −3.91, *p* < 0.001), with rates highest during small pupil, low arousal states (sextile 1 mean ± SEM: 0.042 ± 0.003, sextile 6: 0.033 ± 0.003; Fig. 3e). This effect was significantly greater than that in posterior hippocampus (anterior – posterior slope = –0.002 ± 0.0008 Hz, t(593) = –2.30, *p* = 0.022), where ripple rate did not significantly vary with pupil sextile (posterior hippocampus slope = −0.0002 ± 0.0006 Hz, t(593) = −0.30, *p* = 0.763; sextile 1 mean ± SEM: 0.030 ± 0.003, sextile 6: 0.029 ± 0.003; Fig. 3e). Interestingly, this anterior-posterior asymmetry was driven by elevated ripple rates in anterior hippocampus during the lowest arousal (smallest pupil) states, paralleling the pattern observed in the sleep data (anterior–posterior at sextile 1: t(254) = 3.67, p < 0.001; at sextile 6: t(254) = 0.85, *p* = 0.396; Fig. 3e). Ripple events in anterior hippocampus also showed a pronounced PRW that varied across pupil sextiles and was greatest in magnitude during small pupil, low arousal states (slope (small to large pupil) = 6.90 ± 2.06 µV, t(566) = 3.35, *p* < 0.001; Fig. 3d), further replicating trends observed in the sleep-stage analyses. PRWs were larger in anterior than posterior hippocampus (region: 43.35 ± 16.20 µV, t(364) = 2.68, *p* = 0.007) and posterior hippocampus PRWs did not significantly vary across pupil sextiles (slope = 2.36 ± 2.50 µV, t(569) = 0.94, *p* = 0.347), although the interaction between hippocampal region and pupil sextile was not significant (anterior – posterior slope: 4.54 ± 3.24 µV, t(568) = 1.40, *p* = 0.162).

Beyond the hippocampus, ripple-like events detected in the amygdala during wake had no significant relationship to pupil sextile (mixed effects model with region as hippocampus or amygdala; amygdala: slope (small to large pupil) = 0.0005 ± 0.0004 Hz, t(978) = 1.01, *p* = 0.313; Fig. 3f). This effect was significantly different than that observed in the entire hippocampus (hippocampus – amygdala slope = –0.0017 ± 0.0006 Hz, t(978) = –2.90, *p* = 0.004), with overall event rates also lower in the amygdala (region: –0.0148 ±0.0027 Hz, t(710) = –5.47, *p* < 0.001). Additionally, there was no difference in amygdala PRWs across pupil sextiles (slope = −2.11 ± 1.81 µV, t(928) = −1.16, *p* = 0.245; Fig. 3f), differing significantly from the entire hippocampus (hippocampus – amygdala slope = 7.18 ± 2.32 µV, t(925) = 3.10, *p* = 0.002), and PRWs were smaller in the amygdala than in the hippocampus overall (region: slope = 33.09 ± 11.22 µV, t(656) = 2.95, *p* = 0.003). Results from analyses of the lateral temporal cortex again recapitulated those observed in the amygdala—event rates showed no variation with pupil-linked arousal states (slope = –0.0004 ± 0.0004 Hz, t(1367) = –0.94, *p* = 0.346; Fig. S5)—further suggesting the specificity of hippocampal-arousal modulation. While the fixation task and data processing methods controlled for eye-movements and blinks, we further confirmed such periods did not bias or drive observed effects (Fig. S3).

To better understand the observed differences between anterior and posterior hippocampus, we next investigated how this pupil-ripple modulation varied continuously along the hippocampal long axis (AP position). Like in the sleep stage analyses, for each electrode we calculated a selectivity index, but this time to small pupil (low arousal) states, where small pupil included sextiles 1 and 2 and large pupil included sextiles 5 and 6 (Fig. 3g; see Methods). This index quantified the degree to which ripple rates increased during small pupil states compared to large pupil states; positive and negative values reflected greater ripple rates during small pupil (low arousal) and large pupil (high arousal) states, respectively. Like in the sleep data, SI values visually appeared to cluster anteriorly (Fig 3g). While AP position did not significantly predict SI values (slope (anterior to posterior) = −.186 ± 0.172, t(109.89) = −1.09, *p* = 0.280), in a more robust model predicting ripple rate as a function of an interaction between all pupil sextiles (1-6) and continuous AP position, the pupil sextile × AP position interaction was significant: pupil-ripple modulation was strongest in the most anterior hippocampal electrodes and decreased towards posterior hippocampus (pupil sextile × AP position slope = 0.0046 ± 0.0022, t(593) = 2.04, *p* = 0.042). Altogether, these results demonstrate that during wake, where both ripple rates and arousal dips are generally attenuated compared to sleep, hippocampal ripple events still occur more predominantly during periods of low arousal, mirroring observations from sleep-stage analyses in both anatomical distribution and ripple morphology.

### Hippocampal ripple rate increased during slow heart rate states in both sleep and wake

Consistent with our prediction, hippocampal ripple rates were modulated by arousal state across sleep and wake: ripple rates were highest during low arousal stages of sleep (i.e., NREM) and wake (i.e., small pupil). Across these analyses, sleep stages and pupillometry emerged as complementary measures, with sleep stages capturing large arousal changes in sleep and pupillometry tracking more subtle and transient arousal fluctuations in wake. However, the use of different metrics across sleep and wake does limit direct comparison between states. Since pupillometry was not feasible during overnight recordings in neurosurgical patients, we sought an additional arousal metric that was continuously measurable across both sleep and wake to bridge findings across these states. To this end, we focused on heart rate as a broad physiological arousal index that could be tracked equally across sleep and wake. Heart rate is known to track arousal across sleep and wake states and has been observed to co-vary with both sleep stages (Somers et al., 1993) and pupil size (Bradley et al., 2008; Carro-Domínguez et al., 2025; Y.-H. Chang et al., 2025; Strauch et al., 2022). To measure heart rate, we recorded electrocardiogram (EKG) during both the overnight sleep recordings and the wake fixation task for each participant (see Methods; Table S2). From each recording, we extracted a continuous RR interval timeseries—the time between successive EKG R-peaks—where short intervals reflect fast heart rate (high arousal) and long intervals reflect slow heart rate (low arousal; see Methods). Paralleling the pupil analysis, ripple rates were calculated within RR interval sextiles, with sextile 1 corresponding to small RR intervals (fast heart rate; high arousal) and sextile 6 to large RR intervals (slow heart rate; low arousal). We then examined the relationship between RR interval sextile and ripple rate across sleep and wake.

We first characterized the relationship between EKG RR intervals and ripple events during sleep. Following previous literature, RR intervals exhibited large fluctuations across sleep, co-varying with sleep stages (χ²(2) = 58.87, *p* < 0.001): mean RR interval was largest (heart rate was slowest) during NREM sleep (1.008 ± 0.033 s) and smallest during wake (0.913 ± 0.033 s; all pairwise comparisons significant, *p* < 0.005, Tukey corrected; Cajochen et al., 1994; Snyder et al., 1964; Fig. 4a, b). Across the full sleep recordings, hippocampal ripple rate closely tracked RR interval sextiles with rates highest during large RR interval (slow heart rate, low arousal) states and lowest during small RR interval (fast heart rate, high arousal) states (mixed-effects model with predictors of RR sextiles and hippocampal region; main effect of RR sextile: slope (6−1, large to small interval) = −0.0041 ± 0.0002 Hz, t(575) = −19.51, *p* < 0.001; sextile 6: 0.046 ± 0.002 Hz, sextile 1 mean ± SEM: 0.035 ± 0.002 Hz; Fig. 4d), mirroring results from sleep-stage and pupil analyses. RR interval sextile predicted ripple rate in both anterior (slope = −0.0041 ± 0.0002 Hz, t(577) = −19.47, *p* < 0.001) and posterior hippocampus (slope = −0.0007 ± 0.0002 Hz, t(577) = −2.72, *p* = 0.007). Paralleling patterns observed in the sleep and pupil data, this effect was greater in anterior than posterior hippocampus (anterior – posterior = −0.0034 ± 0.0003 Hz, t(577) = −10.65, *p* < 0.001) with greater rates in anterior than posterior during large RR interval, slow heart rate states (anterior – posterior at sextile 6 = 0.019 ± 0.003 Hz, t(128) = 7.30, *p* < 0.001; anterior – posterior at sextile 1 = 0.002 ± 0.003 Hz, t(128) = 0.93, *p* = 0.356; Fig. 4d). This effect additionally varied continuously along the hippocampal long axis: ripple rate increases during slow heart rate states were largest in the most anterior electrodes and decreased toward the posterior hippocampus, measured both with an RR interval sextile selectivity index (SI slope (anterior to posterior) = −0.31 ± 0.057, t(106.34) = −5.48, *p* < 0.001; Fig. S4a) and an anatomical position × RR interval sextile interaction in a mixed-effects model predicting ripple rate (anatomical position × RR interval sextile slope = : 0.0087 ± 0.0009, t(575) = 9.27, *p* < 0.001; Fig. S4a). Finally, like in the sleep-stage and pupil analyses, PRWs in anterior, but not posterior, hippocampus varied across RR interval sextiles and were greatest in magnitude during large RR interval states (slow heart rate) and smallest in magnitude during small RR interval states (fast heart rate; anterior hippocampus slope (6−1, large to small interval) = 2.89 ± 0.43 µV, t(577) = 6.81, *p* < 0.001; posterior hippocampus slope = −0.0542 ± 0.493 µV, t(577) = −0.11, *p* = 0.913; Fig. 4c).

**Figure 4.**
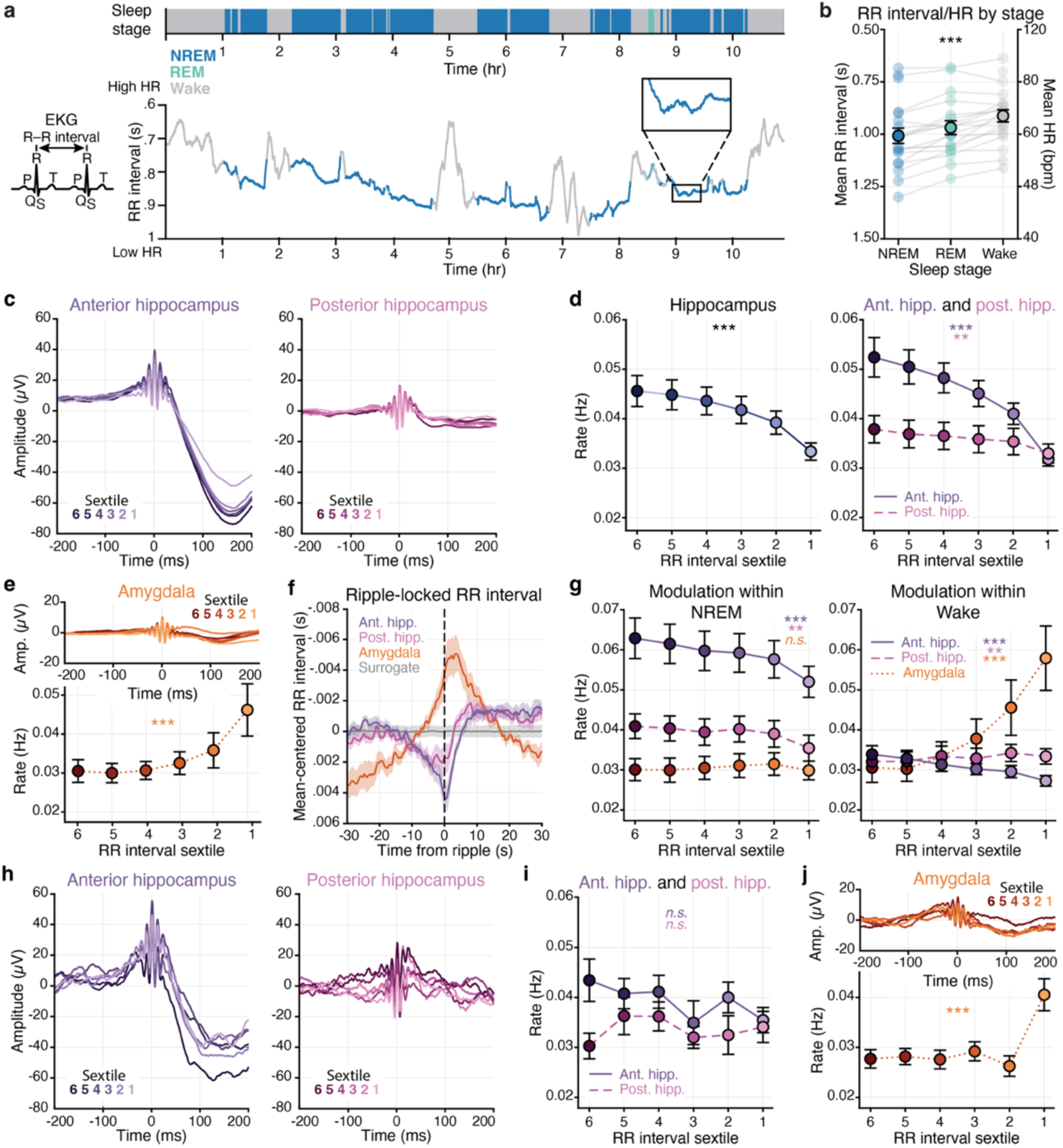
Ripple rate across heart rate states. **a.** EKG RR interval across a representative sleep night (P20). Top bar: sleep stage over time (NREM: dark blue; REM: green; Wake: grey). Y-axis is inverted (higher = shorter RR interval = faster heart rate). **b.** Mean RR interval and heart rate per sleep stage across participants. **c.** Group mean ripple-triggered voltage traces per RR interval sextile during sleep, for anterior hippocampus (left) and posterior hippocampus (right). Line color indicates sextile (dark to light: sextile 6-1, low to high arousal). **d.** Group mean ripple rate across RR interval sextiles during sleep for all hippocampal electrodes (right), and separately for anterior hippocampus (solid) and posterior hippocampus (dashed) (left). **e.** Group mean ripple-triggered voltage traces (top) and ripple rate (bottom) across RR interval sextiles during sleep for amygdala electrodes. **f.** Ripple-locked RR intervals during sleep for anterior hippocampus (purple), posterior hippocampus (pink), and amygdala (orange). Solid lines: mean across participants; shading: ±SEM. Surrogate mean for anterior hippocampus (permutations = 1000) is shown in grey with shading for ±2 surrogate mean SEM. **g.** Group mean ripple rate across RR interval sextiles during NREM (left) and wake (right) periods within sleep recordings, with sextiles calculated within each state, for anterior hippocampus (purple, solid), posterior hippocampus (pink, dashed), and amygdala (orange, dotted). **h., i., j.,** As in c, d, e for the wake fixation task, respectively. Throughout, solid circles: group mean; error bars: ±SEM across group; transparent dots and lines: individual participants; significance of main effect of RR interval sextile: \*\*\**p* < 0.001, \*\**p* < 0.01.

In contrast to the hippocampus, amygdala ripple-like events increased in rate during small RR interval sextile, fast heart rate states (mixed-effects model with region as hippocampus or amygdala; amygdala slope = 0.0025 ± 0.0003 Hz, t(942) = 9.44, *p* < 0.001; hippocampus – amygdala = −0.0051 ± 0.0003 Hz, t(942) = −15.03, *p* < 0.001; Fig. 4e). Additionally, there was no difference in amygdala PRWs across RR interval sextiles (slope: = 0.445 ± 0.391 µV, t(942) = 1.14, *p* = 0.256; hippocampus – amygdala = 1.19 ± 0.5 µV, t(942) = 2.39, *p* = 0.017; Fig. 4e). Results from analyses of the lateral temporal cortex again closely mirrored those observed in the amygdala (Fig. S5).

Leveraging the continuous nature of the RR interval timeseries, we next asked what the temporal dynamics were of this RR interval-hippocampal ripple modulation. To do so, we examined RR interval values locked to ripple event occurrence separately for anterior hippocampus, posterior hippocampus, and amygdala (Fig. 4f). Heart rate was at a minimum (RR interval maximum) at the time of anterior hippocampal ripple events: the RR time course locked to anterior hippocampal ripple events showed an event related modulation that was significantly different than one locked to random events (i.e., surrogate; two-sample KS test vs. surrogate, D = 0.509 ± 0.019, significant in all participants, *p* < 0.001), with heart rate reaching its minimum at the time of the ripple event (RR interval maximum of 0.0044 ± 0.0009 s at 0.36 s). For RR time courses locked to posterior hippocampus ripple events, heart rate similarly reached a minimum around the time of a ripple event (RR interval maximum of 0.0023 ± 0.0007 s at –4.44 s; D = 0.508 ± 0.018, significant in all participants, *p* < 0.001). In contrast, the RR time course locked to amygdala events demonstrated that heart rate reached a maximum around the time of an amygdala event (RR interval minimum of –0.0051 ± 0.0012 s at 3.58 s; D = 0.512 ± 0.025, significant in all participants, *p* < 0.001). Together, these analyses reveal that hippocampal and amygdala ripple events are embedded within opposing heart rate states in sleep unfolding over seconds to tens of seconds—hippocampal ripples during low heart rate and amygdala events during high heart rate—highlighting the regional specificity of this effect and raising the possibility that low arousal states do not merely correlate with, but may actively structure, hippocampal ripple genesis.

While heart rate exhibited large fluctuations across sleep that tracked sleep stages (i.e., increased during wake, decreased during NREM; Fig. 4a, b), heart rate also appeared to vary within sleep stages–fluctuating within NREM and wake themselves, akin to recent reports of sleep microstates (H. Chang et al., 2025; Fig. 4a). We therefore examined the RR interval-ripple relationship within NREM and wake periods of the overnight sleep recordings separately, asking whether ripple rates tracked the subtler, more transient arousal fluctuations that occur within a given stage (Fig. 4a, g). This analysis also allowed us to isolate heart rate-ripple relationships separate from any correlations between heart rate and sleep stages. We thus computed RR interval sextiles using only data from each stage (NREM, wake) and calculated event rates within these stage-specific sextiles for the anterior hippocampus, posterior hippocampus, and amygdala. Within NREM itself, both anterior and posterior hippocampus ripple rate increased during large RR interval, slow heart rate states (anterior hippocampus slope (6-1, large to small interval) = −0.0020 ± 0.0002 Hz, t(577) = −10.24, *p* < 0.001); posterior hippocampus slope = −0.0007 ± 0.0002 Hz, t(577) = −3.13, *p* = 0.002), and this effect was greater in anterior than posterior hippocampus (anterior – posterior = −0.0013 ± 0.0003 Hz, t(577) = −4.31, *p* < 0.001; Fig. 4g). Amygdala ripple-like event rate did not vary with RR interval sextiles within NREM (slope = 0.0002 ± 0.0002 Hz, t(942) = 1.13, *p* = 0.260), differing significantly from the entire hippocampus (hippocampus – amygdala = −0.0016 ± 0.0002 Hz, t(942) = −7.77, *p* < 0.001; Fig. 4g; see Fig. S4d for RR interval SI within NREM). Within the wake periods of the overnight sleep recordings, anterior hippocampus ripple rate increased during large RR interval, slow heart rate states (slope = −0.0012 ± 0.0001 Hz, t(577) = −8.11, *p* < 0.001). In posterior hippocampus, ripple rates increased during small RR interval, fast heart rate states−the reverse of what was observed in anterior hippocampus (slope = 0.0004 ± 0.0002 Hz, t(577) = 2.70, *p* = 0.0072; anterior – posterior: −0.0016 ± 0.0002 Hz, t(577) = −7.33, *p* < 0.001; Fig. 4g). In the amygdala, ripple-like event rate increased greatly during small RR interval, fast heart rate states (slope = 0.0053 ± 0.0003 Hz, t(942) = 20.81, *p* < 0.001), the opposite effect of the entire hippocampus (hippocampus − amygdala: −0.0058 ± 0.0003 Hz, t(942) = −17.73, *p* < 0.001) and a greater increase than the posterior hippocampus (posterior hippocampus − amygdala: −0.0049 ± 0.0004 Hz, t(943) = −12.16, *p* < 0.001; see Fig. S4f for RR interval SI within NREM and wake). In addition to the wake periods of the sleep recordings, we also examined the relationship between RR interval sextile and ripple rate in the wake fixation task. Though trends largely mirrored those observed during wake periods of the sleep recordings, effects were weaker and nonsignificant in the hippocampus, likely reflecting reduced statistical power given the minutes-long fixation blocks compared to the hours-long sleep recordings (e.g., slope of hippocampal ripple rate across RR interval sextile (6-1, large to small interval) = −0.0007 ± 0.0004 Hz, t(595) = −1.906, *p* = 0.057; Fig. 4h-j; see supplemental results). Altogether, these results demonstrate that heart rate based arousal fluctuations predict hippocampal ripple genesis and morphology both across and within sleep stages, indicating that this relationship is not merely a byproduct of sleep stage transitions but is instead continuous and extending into wakefulness. These patterns, and their regional specificity, strikingly mirrored those observed when arousal was indexed via sleep staging and pupil size, underscoring that low arousal selectively favors hippocampal ripple genesis.

### Low arousal states promoted hippocampal ripple genesis across measures

Our goal was to examine the impacts of arousal on hippocampal ripple genesis, as we hypothesized that arousal may provide a unifying account for ripple event occurrence across both sleep and wake states. As predicted, low arousal states consistently promoted higher hippocampal ripple rates across sleep and wake, during both large and small fluctuations in arousal, and across multiple measures of arousal physiology (Fig. 5a). At the broadest level of arousal variation, hippocampal ripple rates were maximal during NREM sleep and minimal during wake. This relationship extended beyond discrete stage boundaries, with ripple rate increasing during low heart rate moments of sleep. This arousal-ripple coupling persisted with smaller arousal variations: ripple rates increased during low heart rate moments within NREM and wake stages individually, but with reduced magnitude, consistent with the narrower arousal ranges sampled within any single stage. Similarly, during the wake fixation task, despite generally lower ripple rates and a smaller arousal range than that spanned across sleep, pupil size predicted ripple rate, demonstrating that the pronounced arousal-ripple relationship during sleep extends to wake. Strikingly, the magnitude of the wake pupil effect approached that of the sleep heart rate effect, despite the smaller arousal range sampled during wake—suggesting that ripple generation may be particularly coupled to the arousal systems reflected by pupil diameter. Across all arousal measures, the effect was consistently greater in anterior compared to posterior hippocampus. In contrast, amygdala ripple-like events demonstrated the opposite relationship to arousal state, with rates highest during the wake periods in the sleep recordings and strikingly increased during moments of high heart rate within these wake periods—reinforcing the regional specificity of arousal-ripple modulation.

**Figure 5.**
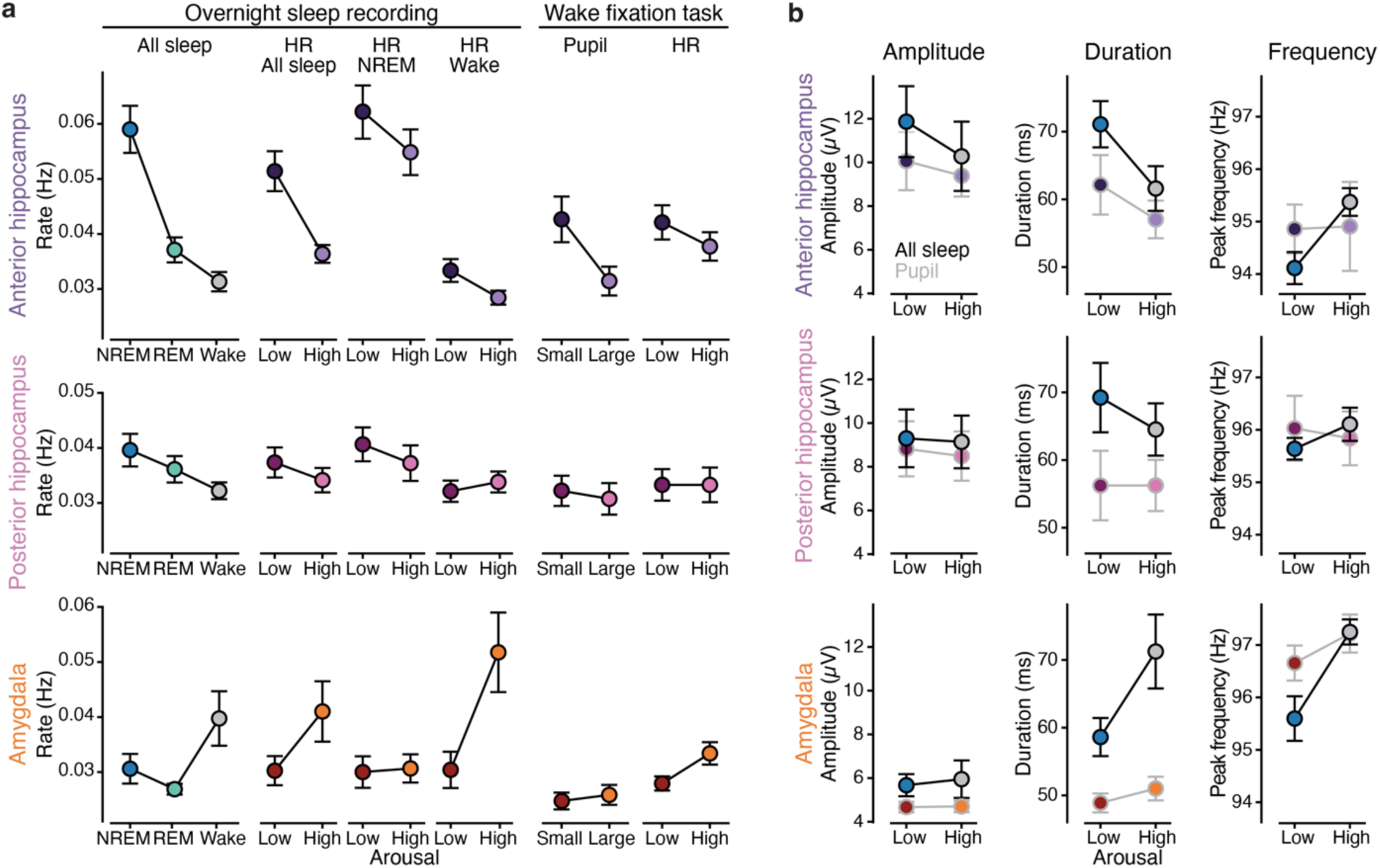
Summary of ripple features across high and low arousal states. **a.** Group mean ripple rate across sleep stage, heart rate, and pupil arousal states, separately for anterior hippocampus (top), posterior hippocampus (middle), and amygdala electrodes (bottom). Heart rate states: low = mean of RR interval sextiles 5 and 6 (slow heart rate), high = mean of sextiles 1 and 2. Pupil states: small = mean of pupil sextiles 1 and 2, large = mean of sextiles 5 and 6. **b.** Group mean ripple amplitude (left), duration (middle), and peak frequency (right), separately for anterior hippocampus (top), posterior hippocampus (middle), and amygdala electrodes (bottom), for low and high arousal states across sleep (sleep stages; black line) and the wake fixation task (pupil states; grey line). Sleep: low arousal = NREM, high arousal = Wake. Wake fixation task: low arousal = small pupil (mean of pupil sextiles 1 and 2), high = large pupil (mean of sextiles 5 and 6). Throughout, solid circles: group mean; error bars: ±SEM across group.

Beyond rate, ripple events vary in morphological features—amplitude, duration, peak frequency—and recent evidence has suggested that these attributes may inform the strength or efficacy of the event (Y. Y. Chen et al., 2021; Fernández-Ruiz et al., 2019; Ngo et al., 2020; Ramirez-Villegas et al., 2015; Robinson et al., 2026; Sebastian et al., 2023; Tong et al., 2021). We characterized these ripple attributes to examine if they were likewise shaped by arousal state (H. Chang et al., 2025). Overall, hippocampal ripple amplitude and duration demonstrated similar arousal modulations as rate in both sleep and wake recordings, with effects strongest in anterior hippocampus: consistent with prior work, anterior hippocampus ripple event amplitude and duration increased during large arousal drops across sleep stages (i.e., NREM; effect of sleep stage on duration: F(2, Inf) = 184.26, *p* < 0.001; on amplitude: F(2, Inf) = 7224.54, *p* < 0.001; Buzsáki, 2015; Y. Y. Chen et al., 2021; Fig. 5b); extending this, for anterior hippocampus both measures also tracked subtle arousal dips during wake indexed by pupillometry (effect of pupil sextile on duration: z = −2.00, *p* = 0.045; on amplitude: z = −2.57, *p* = 0.010; H. Chang et al., 2025; Fig. 5b). Similar but attenuated trends were observed for posterior hippocampus ripple amplitude and duration whereas events detected in the amygdala showed a different pattern of modulation (e.g., largest ripple amplitude and duration during wake periods of sleep recordings; Supplemental results). On average, in mixed-effects models where region represented all three anatomical divisions (anterior hippocampus, posterior hippocampus, or amygdala), ripple event amplitude and duration were overall greatest in anterior hippocampus, attenuated in posterior hippocampus, and substantially smaller in the amygdala in both sleep (main effect of region on duration: χ²(2) = 16.64, *p* < 0.001; on amplitude: χ²(2) = 61.368, *p* < 0.001) and the wake fixation task (main effect of region on duration: χ²(2) = 13.98, *p* < 0.001; on amplitude: χ²(2) = 57.80, *p* < 0.001; see Supplement for peak frequency), underscoring regional differences in ripple event features across MTL structures and within the hippocampus itself (Fig. 5b). In addition to these more commonly analyzed ripple attributes, we characterized the post-ripple wave (PRW)—a morphological feature that, to our knowledge, has not previously been examined in relation to any arousal metric—and found that anterior hippocampal PRW-amplitude similarly tracked arousal across all metrics, growing in magnitude as arousal decreased (Fig. 2c, 3d, 4c). Collectively, the convergence of effects across arousal metrics and ripple event features highlights the robustness of the arousal-dependent hippocampal ripple modulation.

## Discussion

Observations of hippocampal ripples during cognitive tasks in humans challenge the established coupling between ripples, replay, and offline states of inactivity from rodents. Here, we propose that fluctuations in arousal could offer a parsimonious account of ripple genesis across both sleep and active wake states, one that may aid in reconciling the human and rodent ripple literatures. Using invasive intracranial recordings in humans during both sleep and wake, combined with sleep staging, pupillometry, and heart rate measurements of arousal, we show that hippocampal ripples occur predominantly during low arousal states regardless of whether an individual is asleep or awake: ripples preferentially occurred during NREM stages in sleep, small pupil states in wake, and low heart rate states throughout sleep and wake. This robust arousal-ripple relationship was anatomically specific, with the strongest coupling observed in anterior hippocampus, while amygdala ripple-like events showed the opposite pattern, increasing during high arousal states. Together, these findings favor an important role for arousal in shaping when hippocampal ripples occur.

Across three measures of arousal physiology—sleep stages, pupil diameter, and heart rate—hippocampal ripples occurred predominantly during moments of low arousal in both sleep and wake. These results replicate and extend recent rodent findings that reductions in pupil-linked arousal (i.e., pupil constriction) are associated with ripple and replay occurrence during sleep (H. Chang et al., 2025) and ripple occurrence during wake (Farrell et al., 2024; McGinley, David, et al., 2015; Nguyen et al., 2024). Here, we demonstrate the same pupil-ripple relationship in humans during wake and additionally reveal the arousal-ripple relationship using heart rate measures across both sleep and wake. Our findings also align with recent work identifying causal mechanisms that link arousal physiology to hippocampal ripple genesis: both endogenous increases in acetylcholine (ACh) and optogenetic stimulation of medial septal cholinergic neurons were found to suppress hippocampal ripple genesis in rodents, while drops in ACh coincided with ripple occurrence (Vandecasteele et al., 2014; Y. Zhang et al., 2021, 2024). Importantly, ACh has been shown to co-vary with both sleep stages (i.e., low ACh during NREM, high during REM and wake; Marrosu et al., 1995) and pupil diameter (i.e., low ACh during small pupil; Reimer et al., 2016), suggesting that fluctuations in cholinergic tone may be a key mechanism underlying the arousal-ripple relationship we observe. However, ACh is not the only neuromodulator implicated in both pupil and sleep dynamics. Noradrenaline (NA) has also been associated with changes in arousal and pupil diameter (Reimer et al., 2016) as well as with sleep spindle occurrence (Aston-Jones & Bloom, 1981; Kjaerby et al., 2022; Osorio-Forero et al., 2021), which is notable given that ripple-spindle coordination is thought to reflect the hippocampal-cortical communication necessary for memory consolidation (Buzsáki, 2015; Sirota et al., 2003; Staresina et al., 2015). However, the evidence for a direct role of noradrenaline in ripple genesis is mixed (Ul Haq et al., 2012), although there is growing evidence that activity in the locus coeruleus, the brain’s primary noradrenergic nucleus, influences ripple-spindle coupling during sleep (Novitskaya et al., 2016; Swift et al., 2018; Yang & Eschenko, 2025). Together, the convergence of our results in humans with the causal rodent literature suggests that cholinergic arousal in particular, but perhaps in coordination with noradrenergic arousal, is a key determinant of the low arousal states that promote hippocampal ripple genesis.

The coupling between arousal state and ripple occurrence that we observed was not uniform within the hippocampus or across medial temporal lobe structures (i.e., hippocampus and amygdala). Within the hippocampus, arousal-ripple coupling was consistently stronger in anterior than posterior hippocampus, with ripple attributes (e.g., amplitude, duration, etc.) also differing between anterior and posterior hippocampus. The anatomical heterogeneity of ripple events, and the arousal-ripple relationship, aligns with rodent work reporting differences in ripple attributes between dorsal and ventral hippocampus (Buzsáki, 2015; Kouvaros & Papatheodoropoulos, 2017; Sosa et al., 2020; Tingley et al., 2021), recent human work documenting differences in ripple activity between anterior and posterior hippocampus (De Filippo & Schmitz, 2023; Staresina et al., 2023; To et al., 2025; J. Zhang et al., 2025), and a broader literature emphasizing functional and structural differences across the hippocampal long axis (Moser & Moser, 1998; Poppenk et al., 2013; Strange et al., 2014). Critically, in our sleep recordings, while anterior hippocampal ripple rate was significantly elevated compared to posterior hippocampus in NREM, rates of the two regions were matched in wake and REM. The matched rates we observed in wake and REM suggest that the observed regional differences are unlikely to reflect measurement sensitivity, and instead point to potential differences in ripple mechanism, propagation, or function along the hippocampal long axis (Poppenk et al., 2013; J. Zhang et al., 2025). Given recent work in humans examining ripple activity beyond the hippocampus—in other MTL and cortical regions—we examined whether arousal states similarly modulated events detected in the amygdala and lateral temporal cortex. Compared to the hippocampus, events in both the amygdala and cortex showed a strikingly opposite relationship to arousal, with rates being greater during high arousal states. The similarity between the patterns observed in the amygdala and cortex suggests that these effects are not anatomically specific and together may reflect a broader relationship between non-hippocampal ripple-like activity and arousal (Harris & Thiele, 2011; S.-H. Lee & Dan, 2012; McCormick et al., 2020; McGinley, David, et al., 2015). The divergence of the amygdala/cortex pattern from that observed in the hippocampus raises the possibility that ripple-band events detected in non-hippocampal regions may reflect mechanistically distinct phenomena from canonical hippocampal ripples, and thus we refer to them as “ripple-like” throughout (A. A. Liu et al., 2022; Reithler et al., 2025; van Schalkwijk & Helfrich, 2026).

Beyond ripple rate, the post-ripple wave (PRW; also referred to as post-ripple hyperpolarization, post-ripple refractory period; Buzsáki, 2015; Hussin et al., 2020), was found to be sensitive to both arousal state and anatomical location. This attribute is notable because, unlike in rodents, hippocampal ripples detected in humans and non-human primates (NHP) do not consistently exhibit a sharp wave component preceding the ripple oscillation (Jiang et al., 2020; A. A. Liu et al., 2022; Maslarova et al., 2025; Reithler et al., 2025; van Schalkwijk & Helfrich, 2026), however, the PRW appears to be reliably observed across human and NHP recordings. Interestingly, in NHPs, larger PRWs were observed in quiescent states than during task epochs, mirroring observations of larger PRWs during low than high arousal states in humans in our study and emphasizing the potential similarity of this ripple attribute across primate species (Hussin et al., 2020). Previous work suggested the PRW reflects inhibition following a ripple, noting that the prominence of the PRW scaled with the level of activity during the ripple (Buzsáki, 2015; Hussin et al., 2020). Together with this account, our finding that the PRW is sensitive to both arousal and anatomy underscores that it may reflect a physiologically meaningful aspect of ripple genesis or propagation in the primate hippocampus.

Recently, a growing number of human intracranial studies have reported hippocampal ripples during active cognitive task performance. For example, these studies have observed putative ripples in humans during tasks such as visual/perceptual memory (Y. Y. Chen et al., 2021; J. Liu et al., 2026; L. Wang et al., 2026), episodic memory (Norman et al., 2019, 2021; Sakon et al., 2024; Sakon & Kahana, 2022; Vaz et al., 2019), and decision making and planning (He et al., 2026; Zhou et al., 2026). Beyond the observation of ripples during active tasks, these findings present several major departures from prior animal work regarding when ripple events occur. Across species, ripple rates are typically maximal during NREM sleep and minimal during active behaviors (Buzsáki, 2015; Joo & Frank, 2018; A. A. Liu et al., 2022). In addition, changes in ripple rates are typically sensitive to slower shifts in behavioral state, rather than rapidly to sensory stimulation (Buzsáki, 2015; Y. Y. Chen et al., 2021; Leonard et al., 2015; Leonard & Hoffman, 2017). However, several human studies report ripple rates during tasks to be equal to or greater than that observed during NREM sleep in humans (i.e =/> 0.2Hz; Jiang et al., 2019, 2020; other studies observe even lower NREM rates; Y. Y. Chen et al., 2021; Dickey et al., 2022; Ngo et al., 2020; Skelin et al., 2021; Staresina et al., 2015, 2023). Furthermore, recent human studies have reported rapid event-related increases in ripple rate in response to stimulus presentation (e.g., J. Liu et al., 2026; L. Wang et al., 2026).

Together, these findings appear difficult to align with prior animal work on when ripples occur. We suggest two key factors may account for this apparent inter-species difference: i) ripple detection methodologies; ii) arousal state modulations during task performance. First, ripple detection methodologies vary across species and research groups. Many human studies identify ripples from transient increases in power within a predefined ripple-frequency band. Importantly, task engagement and stimulus processing can produce broadband increases in high-frequency activity in the hippocampus (Axmacher et al., 2006; Colgin & Moser, 2010; Henin et al., 2019; Lachaux et al., 2012; Sederberg et al., 2007; Watrous et al., 2015), which can elevate ripple-band power without reflecting a discrete ripple oscillation. If not explicitly rejected, such broadband events could inflate waking ripple rates above that of sleep and create apparent stimulus-related increases. For this reason, prior consensus recommendations have emphasized the importance of spectral confirmation methods that verify ripples as discrete oscillatory events (A. A. Liu et al., 2022; van Schalkwijk & Helfrich, 2026). Indeed, as shown in the present study, the use of ripple spectral confirmation results in a distribution of ripple rates that is highest during NREM sleep and attenuated during wake (Y. Y. Chen et al., 2021), as observed in rodents (Buzsáki, 2015; Joo & Frank, 2018). Second, arousal state may help reconcile ripple occurrence across animal and human studies. The state-dependent framework we propose suggests arousal dips may link ripple occurrence across sleep, quiet wakefulness, and active task performance. In humans, ripples observed during cognitive tasks may arise during transient low-arousal moments embedded within otherwise active behavior. Indeed, our findings show that even subtle reductions in arousal during resting fixation are associated with increased ripple occurrence. This provides a more harmonious account of prior observations of ripples across diverse cognitive tasks: such studies may have captured genuine ripple events, but these events may have reflected endogenous arousal fluctuations rather than the cognitive or sensory events used to structure the analysis. Under this framework, task variables influence ripple occurrence only insofar as they modulate arousal, whereas endogenous arousal fluctuations provide the more proximal predictor of when ripples occur during wakefulness. While this may suggest a re-interpretation of ripple occurrence in previous literature, it would not necessitate revising the functional understanding of ripples in studies where cognitive events converge with arousal variations; instead, this framework may offer a mechanistic account of what promotes ripple occurrence in these studies. Finally, anatomical specificity may also help explain discrepancies across the human ripple literature. In addition to hippocampal ripples, some human studies have included ripple events detected across broader MTL or cortical regions. As our amygdala and cortical findings illustrate, non-hippocampal high-frequency events can show arousal relationships that differ from, or even oppose, those observed in the hippocampus. Consistent with this possibility, prior human studies have reported different ripple patterns in hippocampal versus extra-hippocampal MTL and cortical structures (e.g., Sakon et al., 2024; Sakon & Kahana, 2022). Distinguishing hippocampal ripples from non-hippocampal ripple-band events may therefore help resolve inconsistencies between the human and animal literatures, while clarifying the specific functional role of hippocampal ripples.

While our findings establish a robust arousal-ripple relationship in humans, several limitations exist. First, as our wake recordings were performed during a simple fixation task—chosen to optimize for clean pupil measurements without visual transients or eye movements—no behavioral measure of memory was obtained. This leaves open the question of how the ripple-arousal dynamic we observed relates to mnemonic behavior. However, separate literatures have linked ripples to memory replay and behavior (Buzsáki, 2015; Joo & Frank, 2018), and arousal states to memory performance (Aston-Jones & Cohen, 2005; Clewett & Murty, 2019; Hasselmo, 2006; Sara, 2009), with recent rodent work directly connecting pupil-linked arousal to the expression of memory replay and behavior (H. Chang et al., 2025). Thus, while the arousal framework predicts that endogenous arousal fluctuations should remain the primary determinant of ripple occurrence regardless of task, the finding that low arousal states promote ripple genesis suggests that low arousal moments during mnemonic tasks may impact memory replay and subsequent behavior (H. Chang et al., 2025). Second, the clinical depth electrodes used in our human intracranial recordings do not permit sampling from the large numbers of single neurons necessary to robustly characterize spiking sequences and directly measure memory replay. Recent work has demonstrated ripple-locked activity of stimulus-specific neurons in human hippocampus (Kunz et al., 2024) and ripple-locked memory reactivation in macaque hippocampus (Abbaspoor et al., 2025). However, future work leveraging advances in high-density electrode technology, or recordings in non-human primates where denser neural sampling is currently feasible, in combination with arousal measurements will be critical for establishing whether primate ripple events observed during transient moments of low arousal in wakefulness carry the replay of memory content central to consolidation theories (Reithler et al., 2025). Finally, given our proposal that low arousal states, and in particular moments of low cholinergic arousal, may promote ripple genesis, future work could examine human hippocampal ripples during direct perturbations of cholinergic arousal systems (i.e., pharmacological or neuromodulatory approaches) to provide causal evidence for this relationship in humans.

In conclusion, we report a robust relationship between low arousal states and human hippocampal ripple genesis, suggesting that arousal may be a key driver of ripple occurrence across both sleep and wake. These findings align with a growing focus on arousal as a fundamental factor shaping neural dynamics and behavior (Aston-Jones & Cohen, 2005; McCormick et al., 2020; McGinley, David, et al., 2015; Raut et al., 2025; Sara, 2009). Moreover, a substantial literature has demonstrated that arousal specifically influences mnemonic behavior (Aston-Jones & Cohen, 2005; Clewett & Murty, 2019; Sara, 2009), with fluctuations in neuromodulators, like ACh, thought to drive different states of cognitive focus (i.e., external versus internal; Tarder-Stoll et al., 2020) and memory processing (i.e., encoding versus consolidation; Hasselmo, 2006; Hasselmo & McGaughy, 2004) across wakefulness. Our suggestion that fluctuations in arousal may drive ripple occurrence in both wake and sleep, together with this literature, supports a broader view in which ripples and memory consolidation are not behaviorally segregated from other cognitive processes, restricted to sleep alone. Instead, transient dips in arousal could promote the emergence of ripples and consolidation intermixed with ongoing cognition. This dynamic view aligns with growing evidence suggesting that the brain may continuously fluctuate between external (i.e., online; encoding) and internal (i.e., offline; consolidation) states across multiple timescales, spanning the broader behavioral contexts of sleep and wake (Biba et al., 2026; Honey et al., 2018; Verschooren et al., 2026; Wamsley et al., 2023; Wamsley & Summer, 2020), with arousal driving these transitions. Altogether, this work supports a view of memory formation not as a single event, but instead as a process extended across time, one that may be guided, moment to moment, by arousal state.

## Methods

### Human participants

Intracranial recordings were obtained from 20 human participants (10 F/10 M; mean age: 39.75 years, age range: 21–64 years; see Table S4) undergoing invasive monitoring as part of their treatment for refractory epilepsy at the Hospital of the University of Pennsylvania (Philadelphia, Pennsylvania, USA). One participant (P4) underwent two surgeries with distinct electrode implants, yielding 21 implants across these 20 participants. Recordings were performed using stereo-electroencephalography (sEEG) depth electrodes (Ad-Tech Medical Instrument Corp., WI, USA). All experimental procedures were approved by the Institutional Review Board at the University of Pennsylvania Perelman School of Medicine (IRB protocol number: 821778), with participants providing verbal and written consent.

### Experimental Design

We tested the hypothesis that low arousal states are associated with hippocampal ripple genesis across sleep and wake. To do so, each participant underwent intracranial recordings during overnight sleep and a wakeful fixation task with pupillometry, enabling within-participant analyses of hippocampal ripples across the large arousal fluctuations that accompany sleep stage transitions (measured by sleep staging and electrocardiogram, EKG) and the finer fluctuations present during wakefulness (measured by pupillometry and EKG; Fig. 1).

### Fixation Task

To examine how arousal fluctuations during wakefulness influence ripple genesis, we employed a simple resting fixation task with simultaneous pupillometry, EKG, and intracranial recordings. A fixation task was chosen to optimize pupil recordings by limiting confounds from visual transients and eye movements, allowing measurement of endogenous pupil-linked arousal fluctuations during resting wakefulness (T. Liu et al., 2026).

The task was presented on an adjustable monitor (27”, 3840 × 2160; connected to an Alienware laptop m18 R2 running Windows 11) at a viewing distance of 57 cm using MATLAB (R2023a, MathWorks, MA, USA) and Psychtoolbox (3.0.19; Brainard, 1997). To optimize pupil measurements, low level lighting conditions were kept constant across recordings by closing all blinds, turning off all lights and monitors, and then turning on the same dim lighting configuration in all patient rooms (Mathôt & Vilotijević, 2023). To confirm controlled and dim lighting conditions, illuminance, measured next to the participant’s head at eye level with the task monitor on, was recorded at the start of each data collection period (Dr.meter LX1330B Digital Illuminance Light Meter; mean ± SD: 13.43 ± 5.73 lux).

During the fixation task, participants completed a series of trials, each consisting of an eyes-open phase followed by an eyes-closed phase (Fig. 1d). During the eyes-open phase, participants fixated on a central cross for 17 s before being visually prompted to ‘close eyes’. Participants were instructed to minimize blinking during these eyes-open phases. During the eyes-closed phase (3 s), participants would blink or keep their eyes shut until a brief auditory tone was played to signal the return to eyes-open fixation. To motivate participants’ engagement, on 30% of eyes-open phases the fixation cross slowly rotated at a random time and participants were instructed to indicate the direction of this rotation by pressing the left or right arrow key on a handheld keypad. Experiments were broken into task blocks containing either 10 or 15 eyes-open fixation periods, depending on the participant’s comfort to perform extended fixations, with participants completing at least three task blocks. See Table S1 for the number of task blocks and trials per participant.

### Pupil recordings and preprocessing

Simultaneous pupillometry and intracranial recordings in the epilepsy monitoring unit present practical challenges, as patients are typically reclining in bed and wearing large head wraps. To accommodate this setup, we employed a head-mounted pupillometry system and designed custom 3D-printed flexible frames to secure Pupil Labs world and dual eye cameras (Pupil Labs Core decased, 120 Hz world camera, 200 Hz binocular eye cameras; Fig. 1d). This setup allowed us to record eye camera video synchronized to neural signals via ZMQ (ZeroMQ 4.0, matlab-zmq: https://github.com/fagg/matlab-zmq), with continuous measurements of pupil diameter extracted from the eye camera videos. The pupillometry system was calibrated at the onset of each data-collection session using single-point calibration, with additional calibration performed as needed between blocks. Pupil diameter (mm) from both eyes was exported using Pupil Labs Pupil Player (v3.5.7) before being post-processed and analyzed using custom scripts with MATLAB 2022b.

Artifacts and blinks were detected and removed using the Pupil Labs blink detector (Pupil Player v3.5.6), which identifies blinks based on abrupt changes in the confidence of the pupil-ellipse fit. An additional 50 ms was removed before and after each detected event to account for a blink-associated pupillary response (Diamond, 2001; Yoo et al., 2021). The pupil data were then manually inspected to ensure complete blink removal; if any blink-associated response remained, additional samples were removed until the data were fully blink-free. Outlier pupil values that exceeded 2.5 SD from the mean were additionally removed as likely signal artifacts. Trials were excluded from a block if more than 70% of the trial was removed or if the blink rate exceeded 0.5 blinks/second and more than 30% of the trial was removed. If more than 50% of trials in a block were excluded, that block was excluded from analyses.

After blink and artifact removal, data were interpolated using linear interpolation and low-pass filtered below 4 Hz before being z-scored within blocks for final analyses (Krishnamurthy et al., 2017). Data from a single eye were used for analyses—with the right eye used by default, or the left eye if the camera angle yielded a suboptimal recording from the right. Only eyes-open periods of the wake fixation task were included in pupil analyses.

### Electrophysiological recording

Intracranial sEEG data during sleep and the wake fixation task were acquired at a sampling rate of 2 kHz with a bandpass of 0.3–500 Hz (4^th^-order Butterworth filter) using a BlackRock Cerebus system (BlackRock Microsystems, UT, USA). Initial recordings were referenced to a depth electrode contact within the skull or white matter, distant from pathological zones. During the wake fixation task, trial events were tracked using a photodiode sensor (attached to the stimulus monitor) synchronously recorded at 30 kHz. All additional data processing was performed offline.

### Sleep recording and scoring

To examine how large arousal fluctuations during sleep influence ripple genesis, overnight sleep recordings were acquired from all participants. Data were recorded as described in *Electrophysiological recording*, with the experimenter manually starting and stopping the recording.

To examine the relationship between sleep stages (as a metric of arousal) and ripples, sleep scoring was performed (Fig. 1d) for each recording in two steps: automated scoring using Yet Another Spindle Algorithm (YASA; Vallat & Walker, 2021) followed by manual checking by an expert scorer following international standards (Iber et al., 2007). To automatically score sleep, sEEG data from every electrode across the brain were downsampled to 125 Hz and then bandpass filtered between 0.4–30 Hz. Next, the YASA algorithm was applied to classify the data into awake, rapid-eye movement (REM), and non-REM (NREM) sleep stages (N1, N2, N3). YASA computed a sleep score and confidence value for each electrode for 30 s epochs across the night, resulting in a sleep stage array of epochs by electrode. Electrodes whose confidence (0–1) was less than 0.6 were excluded, and epochs were excluded if more than 50% of electrodes were removed. Each remaining epoch was assigned the sleep stage with the highest average confidence across retained electrodes.

All YASA-generated scores were manually checked by an expert scorer (ES) using MATLAB (R2024b) and Danalyzer (https://github.com/ddenis73/danalyzer; Denis et al., 2021) according to standard criteria (Iber et al., 2007). As polysomnography was not available for all participants, manual scoring was performed by visualizing the sEEG data. EKG, EOG, and EMG were used to supplement scoring when available, though stage determinations did not require these signals given the large number of sEEG electrodes available. For nights with an average YASA confidence exceeding 70% (across all sEEG electrodes), transitions between sleep stages were manually checked and corrected. For nights where the average YASA confidence score was below 70%, the entire night was manually reviewed and corrected.

We aimed to collect two nights of sleep data per participant to reduce the impact of night-to-night variability in sleep architecture and confirm recording quality. If a participant had a seizure overnight or did not achieve at least one hour of NREM sleep and one hour of wake, additional nights were recorded until two clean nights were obtained. For one participant (P1), only one night met these quality criteria and was used in analyses. See Tables S2 and S3 for sleep duration details across participants.

### Electrode localization and selection

To identify sEEG electrode contacts within the hippocampus, amygdala, and cortex, a post-operative CT scan was co-registered to a pre-operative T1 anatomical MRI scan for each participant using the YAEL (Your Advanced Electrode Localizer; Z. Wang et al., 2023) module of RAVE (Reproducible Analysis and Visualization of iEEG software; Magnotti et al., 2020), with 3D coordinates for each electrode computed with RAVE. For each participant, electrodes located in the hippocampus and amygdala were manually identified in native space, with visualization performed in RAVE and ITK-SNAP (v3.6.0; Yushkevich et al., 2006). Additionally, for each depth electrode targeting these structures, a white matter contact and the most lateral grey matter contact were identified to serve as the re-reference (described below) and comparison recording site (see Supplement) for hippocampal and amygdala recordings.

Given known functional differences across the long axis of the hippocampus (Moser & Moser, 1998; Poppenk et al., 2013; Strange et al., 2014), we sought to classify hippocampal electrodes as anterior or posterior. To do so, an expert (EMS) manually identified the uncal apex for each participant’s left and right hippocampi, with electrodes anterior to this landmark assigned to anterior hippocampus and those posterior assigned to posterior hippocampus.

We additionally sought to quantify and aggregate each electrode’s relative position along the hippocampal long axis to obtain a fine-grained understanding of ripple activity across this structure. To aggregate hippocampal recording sites across participants with differing anatomy, we (i) generated a hippocampal segmentation in each participants native MRI space, (ii) aligned each hippocampus to the same coordinate frame, and then (iii) projected all electrode coordinates into this normalized space. To generate hippocampal segmentations, each participant’s T1 MRI was nonlinearly registered to the MNI152 template using ANTs (Advanced Normalization Tools; Avants et al., 2009), and the MNI-space anatomical segmentation (aseg atlas) was then warped back into native T1 space using nearest-neighbor interpolation, yielding a participant-specific hippocampal parcellation (using YAEL and RAVE). To align hippocampi across participants, a 3D surface mesh of each hippocampus was reconstructed from the segmentation using an isosurface algorithm with Laplacian smoothing. Principal components analysis (PCA) was applied to the mesh vertices as a mathematical shortcut to rigidly align each hippocampus to an anatomically consistent coordinate frame. Hippocampal meshes were rotated so that their longest dimension, the anterior-posterior axis, was aligned to the RAS coordinate system. Finally, each electrode’s coordinates were projected into this normalized space, and its position along the long axis was expressed as a proportional distance between the most anterior and most posterior extent of the hippocampus, yielding a normalized anterior-to-posterior position (0 = most anterior, 1 = most posterior).

### EKG recording and preprocessing

EKG data were recorded during both overnight sleep and the fixation task as a continuous measure of arousal applicable across states and datasets.

EKG was synchronously recorded with the neural signals using two clinical EKG electrodes, sampled at 2 kHz, with the BlackRock Cerebus system (see “Electrophysiological Recording”). EKG data were preprocessed offline using MATLAB v2024b. Signals were first downsampled to 1 kHz. To remove large artifacts in the signal, a bipolar EKG was computed from the two electrodes. Both individual and bipolar EKG signals were visually inspected; recordings were rejected if EKG R-peaks were not visible or additional large artifacts were present in the signal.

RR intervals, the difference between EKG R-peaks, were used as a continuous measure of heart rate across time. To detect R-peaks, the bipolar EKG signal was bandpass filtered between 2–80 Hz, followed by a notch filter at 60 Hz (if one EKG electrode was rejected, R-peaks were detected using the remaining electrode). R-peaks were amplified using a continuous wavelet transform (Daubechies 3 wavelet, scale = 0.08 s, corresponding to approximate QRS complex duration) prior to detection with the findpeaks function in MATLAB. RR intervals were then calculated as the temporal difference between R-peaks in seconds. RR intervals less than 0.5 s or greater than 1.7 s were deemed artifactual and removed. The remaining RR interval time series was interpolated to 50 Hz using the PCHIP method (Benchekroun et al., 2023; C.-W. Chen et al., 2023; Jacobsen et al., 2026).

EKG was successfully recorded and RR intervals were calculated for all participants. One participant (P16) had unusable EKG during one of two sleep nights, resulting in the exclusion of that night from EKG analyses. One participant had two separate implants and hospital stays (as noted previously, P4). EKG was not recorded during the sleep nights of this participant’s second implant, resulting in the exclusion of two additional nights. In total, 38 nights of sleep were used for EKG analyses, with at least one night contributed by every participant (n = 20; Table S2).

### Quantification and statistical analysis

#### sEEG preprocessing and spectral decomposition

All sEEG signal processing was performed using custom MATLAB scripts (R2022b & R2024b, MathWorks, MA, USA). All identified hippocampal and amygdala electrodes (see *Electrode Localization and Selection*) were notch filtered (60 Hz and harmonics) and re-referenced to a proximal electrode contact within the white matter on the same probe (Y. Y. Chen et al., 2021). Re-referenced signals from each electrode were downsampled to 1 kHz and spectrally decomposed using Morlet wavelets (7 cycles) with center frequencies spaced linearly from 1 to 200 Hz in 1 Hz steps.

#### Ripple detection

After preprocessing, ripple events were identified across regions (hippocampus, amygdala, cortex) in three stages: i) time-domain detection for identifying putative ripple events; ii) frequency-domain assessment for accepting or rejecting ripple events; iii) electrode-wise ripple rejection thresholding for inclusion or exclusion (Y. Y. Chen et al., 2021); following recent consensus methodologies (A. A. Liu et al., 2022).

Time-domain detection: Notch filtered and re-referenced signals were bandpass filtered from 80 to 120 Hz (ripple band) using a 4_th_-order FIR filter. The root mean square (RMS) of the bandpass-filtered signal was calculated and smoothed using a 20 ms window. Ripple events were identified as periods with ripple band RMS amplitude between 2.5 and 9 standard deviations above the mean. Detected ripple events with a duration shorter than 38 ms (3 cycles at 80 Hz) were rejected.

Frequency-domain assessment: The spectral amplitude distribution (1–200 Hz) for each detected ripple event was computed by averaging the normalized instantaneous amplitude between the onset and offset of the ripple event (using spectrally decomposed data). Spectral amplitude was normalized to percent change from baseline by applying a correction at each frequency based on the mean amplitude across the entire recording for a given electrode and frequency. For the spectral distribution of each detected ripple event, spectral peaks were identified using the *findpeaks* function in MATLAB, and events were rejected if: (1) there was more than one peak in the ripple band; (2) the most prominent and highest peak fell outside the ripple band; (3) the ripple peak width exceeded 3 standard deviations from the mean peak width for that electrode and recording session; or (4) high-frequency activity peaks exceeded 80% of the ripple peak height (following steps from Y. Y. Chen et al., 2021). Any remaining ripples with amplitude exceeding 300 µV were considered artifactual and rejected. This spectral assessment was designed to ensure detected events had a frequency profile consistent with true oscillatory ripples, thereby excluding other high-frequency activity (i.e., broadband gamma and interictal spikes) that may overlap temporally but differ spectrally (Y. Y. Chen et al., 2021; A. A. Liu et al., 2022; Maslarova et al., 2025; van Schalkwijk & Helfrich, 2026).

Electrode-wise ripple rejection thresholding: the ripple rejection rate was calculated for each electrode within each recording block. Electrodes with a rejection rate exceeding 60% were excluded from that recording block, as these likely reflected noisy recordings. Electrodes excluded from 75% or more of recording blocks were removed from all analyses.

### Statistical analysis

To test our hypothesis that low arousal is coupled with increases in hippocampal ripples across states, we quantified the relationship between arousal (sleep stage; pupil size; RR interval) and ripple attributes (rate; amplitude; duration; peak frequency; post-ripple wave). To enable rate-based analyses, arousal metrics were divided into discrete windows. Sleep was split into stages (W, REM, N1, N2, N3), with main analyses focused on comparisons of ripple rate across W (high arousal), REM, and NREM (low arousal; defined as N2 and N3; N1 was excluded due to its transitional nature between wake and sleep). Pupil size and RR intervals were each divided into sextiles within each recording block, with sextiles chosen to align with recent pupil-ripple quantification in the rodent (H. Chang et al., 2025; Fig. 3c). Small pupil sextiles (i.e., sextile 1) corresponded to small pupil and low arousal states, while large pupil sextiles (i.e., sextile 6) corresponded to large pupil and high arousal states. Large RR interval sextiles (i.e., sextile 6) corresponded to slow heart rate and low arousal states, while small RR interval sextiles (i.e., sextile 1) corresponded to fast heart rate and high arousal states. For Fig. 5, pupil sextiles 1 and 2 were grouped to represent a small pupil state (low arousal), and sextiles 5 and 6 to represent a large pupil state (high arousal); RR interval sextiles 5 and 6 were grouped as a low heart rate state (low arousal) and sextiles 1 and 2 as a high heart rate state (high arousal)

#### Mixed-effects analyses of arousal-ripple relationships

Across analyses, the relationships between arousal (pupil size; sleep stage; RR interval) and ripple attributes (rate; amplitude; duration; peak frequency; post-ripple wave) were assessed using linear mixed-effects models (*lme4* package; Bates et al., 2015) in R (R Core Team, 2021). In all models, ripple attributes were predicted by arousal as a fixed effect, and participant and electrode were included as random effects, to accommodate nested data (electrodes) within participants. For analyses of ripple modulation differences between the hippocampus and amygdala, and across the hippocampal long axis, electrode location (hippocampus/amygdala; anterior/posterior; anterior-to-posterior proportional position) was included as a categorical fixed effect with an interaction with arousal.

Mixed-effects models were assessed using a combination of approaches depending on predictor type. For categorical predictors (i.e., sleep stage), the *Anova* function from the *car* package (Fox & Weisberg, 2019; Type II Wald chi-square tests) was used to assess overall significance of each fixed effect, with *p*-values for individual factor levels obtained via *lmerTest* (Kuznetsova et al., 2017) using Satterthwaite’s degrees of freedom method (which provides a conservative adjustment accounting for the complexity of the mixed-effects structure). Post hoc comparisons were performed using the *emmeans* package (Lenth, 2022), with Tukey’s adjusted *p-*values and Kenward-Roger degrees of freedom, to control for multiple comparisons while maintaining statistical power through improved degrees of freedom estimation. For continuous predictors (i.e., pupil sextile, RR interval sextile), significance was assessed from model coefficients via *lmerTest*, providing direct inference on linear effects; where continuous predictors interacted with categorical factors, estimated slopes were obtained using *emtrends* with Kenward-Roger degrees of freedom, enabling interpretation of how continuous effects (e.g., impact of pupil sextile on ripple rate) vary across categorical levels (e.g., anatomical region). For analyses of ripple amplitude and duration, given the large sample size (each ripple an independent sample, versus rate calculations which collapse counts within time bins), degrees of freedom were estimated using the asymptotic (z) approximation, as Kenward-Roger and Satterthwaite corrections converge on this solution at large sample sizes and impose computational costs without meaningful differences in inference. Across models, assumptions of normality, linearity, and homoscedasticity were confirmed using the *resid_panel* function from the *ggResidpanel* package (Goode & Rey, 2022).

#### Ripple-locked RR intervals

To evaluate the relationship between RR intervals and ripple events across time, we examined the ripple-locked RR-interval timeseries (Fig. 4f). RR intervals were extracted in a ±30 s window time-locked to each ripple event, averaged across epochs within each participant, and mean centered, separately for anterior hippocampus, posterior hippocampus, and amygdala (Fig. 4f). A surrogate distribution was constructed for each region by repeating this procedure 1000 times, replacing true ripple event times with an equal number of randomly selected timepoints drawn from the same recording session, yielding a null distribution of 1000 participant-averaged RR interval epochs per region. True and surrogate RR interval distributions were compared using a two-sample Kolmogorov-Smirnov test conducted separately for each participant, with the KS statistic D reported as a measure of effect size.

#### Low arousal selectivity index

To evaluate how arousal-ripple modulation varied across the hippocampal long axis, we computed a selectivity index (SI) for each electrode quantifying the degree to which ripple rates were elevated during low relative to high arousal states: SI = (low arousal rate – high arousal rate) / (low arousal rate + high arousal rate). For sleep stage analyses, ripple rates in NREM and wake were used for low and high arousal rates, respectively. For pupil analyses, ripple rates during small pupil states (mean rate of sextiles 1 & 2) and large pupil states (mean of sextiles 5 & 6) were used for low and high arousal states, respectively. For heart rate analyses, ripple rates during large RR intervals (slow heart rate; mean of sextiles 5 & 6) and small RR intervals (fast heart rate; mean of sextiles 1 & 2) were used for low and high arousal states, respectively. In all cases, positive SI values indicated greater ripple rates during low arousal states and negative values indicated greater ripple rates during high arousal states. Electrodes were plotted by their AP position and colored by their SI value. We statistically evaluated whether SI differed across the long axis for each arousal measurement using mixed effects models predicting SI as a function of continuous AP position, with electrodes nested within participants as intercepts.

## Acknowledgments

The authors thank the patients who participated in this experiment and the staff at the Hospital of the University of Pennsylvania Epilepsy Monitoring Unit for their assistance. The authors also thank Dr. Pin-Chun Chen for helpful discussions on heart rate analyses, and Ilaina Edelstein and Dr. Zhengjia Wang for help with electrode localization.

## Author contributions

Conceptualization, E.M.S., Y.Y.C., A.C.S., B.L.F.; data collection, E.M.S., Y.Y.C., K.A.D., H.I.C; data analysis, E.M.S., Y.Y.C.; resources, K.A.D., H.I.C, B.L.F.; writing, E.M.S., B.L.F., A.C.S.; supervision, B.L.F., A.C.S.

