## Supplemental Information for "Arousal state modulates human hippocampal ripples"

Siefert et al.

### Supplementary Text:

#### Results

***Hippocampal ripple rate was highest in N3 sleep***

The main result of sleep stage-ripple modulation in the hippocampus was replicated in an analysis where all stages were considered individually (N3, N2, N1, REM, Wake; Fig. S2). Briefly, anterior hippocampus ripple rates strongly co-varied with sleep stage: rates were highest in N3 (lowest arousal state), decreased to N2 and N1, and were lowest in wake (highest arousal stage). In posterior hippocampus, ripple rate again co-varied with sleep stage but with attenuated magnitude: rates were highest in N3 and N2 and lowest in N1 and wake. The pattern in the amygdala differed from both the anterior and posterior hippocampus. See Fig. S2 for a more detailed description and statistics.

***Small pupil sextiles predicted high ripple rates even when ripples that co-occurred with blinks were removed***

The main result of pupil-ripple modulation in the hippocampus was replicated in an analysis where ripples that occurred during blinks were removed: To confirm that the inclusion of interpolated pupil values from blink periods did not drive the reported pupil-ripple modulation, we evaluated the pupil-ripple relationship again after excluding any ripples that occurred within 200 ms of a detected blink. Replicating the main analyses, anterior hippocampus ripple rate significantly tracked pupil sextile, decreasing as pupil size increased, while posterior hippocampus and amygdala showed no significant relationship to pupil sextile. See Fig. S3 for a more detailed description and statistics.

***Ripple rate modulation by RR interval sextiles in the wake fixation task***

Ripple rate modulation by RR interval sextiles was analyzed in the wake fixation task as an additional period of wakefulness within which results could be compared to those from pupil-ripple analyses. First, ripple rates were calculated for RR interval sextiles computed across full wake fixation task blocks. Next, mixed-effects models were applied, predicting ripple rate as a function of RR interval sextiles (1–6) and region (anterior vs posterior hippocampus; hippocampus vs amygdala), with electrodes nested within participants as intercepts. The effect of RR interval sextile on hippocampal ripple rate did not reach significance, but there were marginal increases in ripple rate during slow heart rate states in the entire hippocampus and anterior hippocampus (entire hippocampus slope (6-1, large to small interval) = −0.0007 ± 0.0004 Hz, t(595) = −1.906, $p$ = 0.057); anterior hippocampus slope: = −0.0008 ± 0.0004 Hz, t(597) = −1.903, $p$ = 0.058; posterior hippocampus slope: = 0.0002 ± 0.0005 Hz, t(597) = 0.478, $p$ = 0.632; Fig. 4i; see Fig. S4h for visualization of this effect along the hippocampal long axis continuously via SI). Anterior and posterior hippocampus post-ripple waves (PRWs) did not vary significantly with RR interval sextile (anterior hippocampus slope = 0.028 ± 1.57, t(590) = 0.018, $p$ = 0.986; posterior hippocampus slope = 0.523 ± 1.86, t(590) = 0.282, $p$ = 0.778; Fig. 4h). As in the wake periods of the sleep recordings, amygdala ripple-like event rate increased significantly during small RR interval, fast heart rate states (slope = 0.0017 ± 0.0004 Hz, t(982) = 4.423, $p < 0.001$), the inverse of the hippocampal trend (hippocampus − amygdala: −0.0021 ± 0.0005 Hz, t(982) = −4.147, $p < 0.001$; Fig. 4j). There was no difference in amygdala post-event amplitudes across RR interval sextiles (slope = 0.288 ± 1.26, t(971) = 0.228, $p = 0.820$; Fig. 4j).

***Increases in hippocampal ripple rate in low heart rate states followed an anterior-to-posterior gradient***

As with sleep stages and pupil size, the relationship between heart rate (RR interval sextile) and ripple rate across the hippocampal long axis was evaluated. Across sleep, within individual sleep stages (i.e., within NREM, wake), and in the wake fixation task, the largest ripple rate increases in low heart rate states were observed anteriorly, with effects decreasing towards posterior hippocampus. This anterior-to-posterior gradient mirrored that observed in analyses of sleep stages and pupil size. See Fig. S4 for visualization of a heart rate selectivity index across the hippocampal long axis, and a more detailed description and analysis of this result.

***Ripple event attributes across arousal states and anatomical regions***

In addition to rate, ripple attributes of amplitude, duration, and peak frequency were analyzed for their relationships to arousal state. Anterior hippocampus ripple amplitude and duration increased during low arousal sleep stages and pupil states, mirroring observations of rate (see main text; Fig. 5b). Posterior hippocampus ripple amplitude and duration also co-varied with sleep stages (amplitude: F(2, Inf) = 963.77, $p < 0.001$; duration: F(2, Inf) = 32.847, $p < 0.001$) and amplitude tracked pupil size (z = −3.046, $p = 0.002$), although the relationship between duration and pupil size did not reach significance (z = −0.120, $p = 0.904$; Fig. 5b). For the amygdala, amplitude and duration of ripple-like events demonstrated trends that countered the observations in the hippocampus, being greatest during wake (duration: F(2,Inf) = 229.84, $p < 0.001$; amplitude: F(2, Inf) = 148.49, $p < 0.001$) and showing no relationship to pupil size (duration: z = 0.318, $p = 0.750$; amplitude: z = 0.399, $p = 0.690$; Fig. 5b). There were also overall differences in mean amplitude and duration across regions, with anterior hippocampus events having the largest amplitudes and durations and amygdala events having the smallest amplitudes and durations (see main text).

Ripple peak frequency was highest in wake and lowest in NREM in both anterior and posterior hippocampus (anterior: F(2, Inf) = 184.26, $p < 0.001$; posterior: (F(2, Inf) = 32.85, $p < 0.001$), as well as in the amygdala where this effect was especially pronounced (F(2, Inf) = 1712.12, $p < 0.001$). Pupil size did not significantly predict peak frequency in anterior or posterior hippocampus (anterior: z = 1.33, $p = 0.185$; posterior: z = 0.366, $p = 0.714$), or in the amygdala (z = 1.38, $p = 0.169$). Overall, mean peak frequency differed across anatomical regions, with amygdala events demonstrating the highest peak frequencies and anterior hippocampus events the lowest peak frequencies in both sleep (main effect of region: $\chi^2(2) = 34.70$, $p < 0.001$) and the wake fixation task (main effect of region: $\chi^2(2) = 11.48$, $p = 0.003$; Fig. 5b). These results suggest that ripple peak frequency, while sensitive to anatomical region and sleep stages, may not track arousal across behavioral states (e.g., sleep, wake) in the same way as rate, amplitude, duration, and the post-ripple wave. This observation is in line with rodent literature that reported peak frequency to co-vary with amplitude across sleep but diverge during exploratory pauses during wakefulness, suggesting that these features may be driven by partially dissociable mechanisms depending on state (Buzsáki, 2015).

***Ripple-like events detected in lateral temporal cortex displayed a similar relationship to arousal as that from the amygdala***

The relationship between arousal and ripple-like events detected in lateral temporal cortex was evaluated as an additional control to assess the specificity of the amygdala results, and by proxy the hippocampal results. To do so, ripple-like events were detected on the most lateral grey matter contact on each depth electrode targeting hippocampus and amygdala and analyzed for their relationship to arousal states (sleep stages, pupil size, heart rate) in both the sleep and wake fixation task recordings (Fig. S5a, see Methods). Across recordings and arousal states, patterns of event rates in cortex mirrored those in the amygdala and opposed those in the hippocampus, highlighting the specificity of the hippocampal effect and supporting the use of the amygdala as a representative control region (Fig. S5c-d). See Fig. S5 for a more detailed description and statistics.

**Supplementary Figures & Tables:**

**Figure S1**. Sleep stages and durations across participants.
**Figure S2**. Ripple rate across sleep stages—additional participant and stages.
**Figure S3**. Ripple rate across pupil states with blink window.
**Figure S4**. Ripple rate selectivity to RR interval sextiles across sleep and wake.
**Figure S5**. Ripple-like burst activity in lateral temporal cortex across arousal states.

**Table S1**. Experimental data summary.
**Table S2**. Sleep information for individual participants.
**Table S3**. Sleep duration summary across participants.
**Table S4**. Participant demographics.

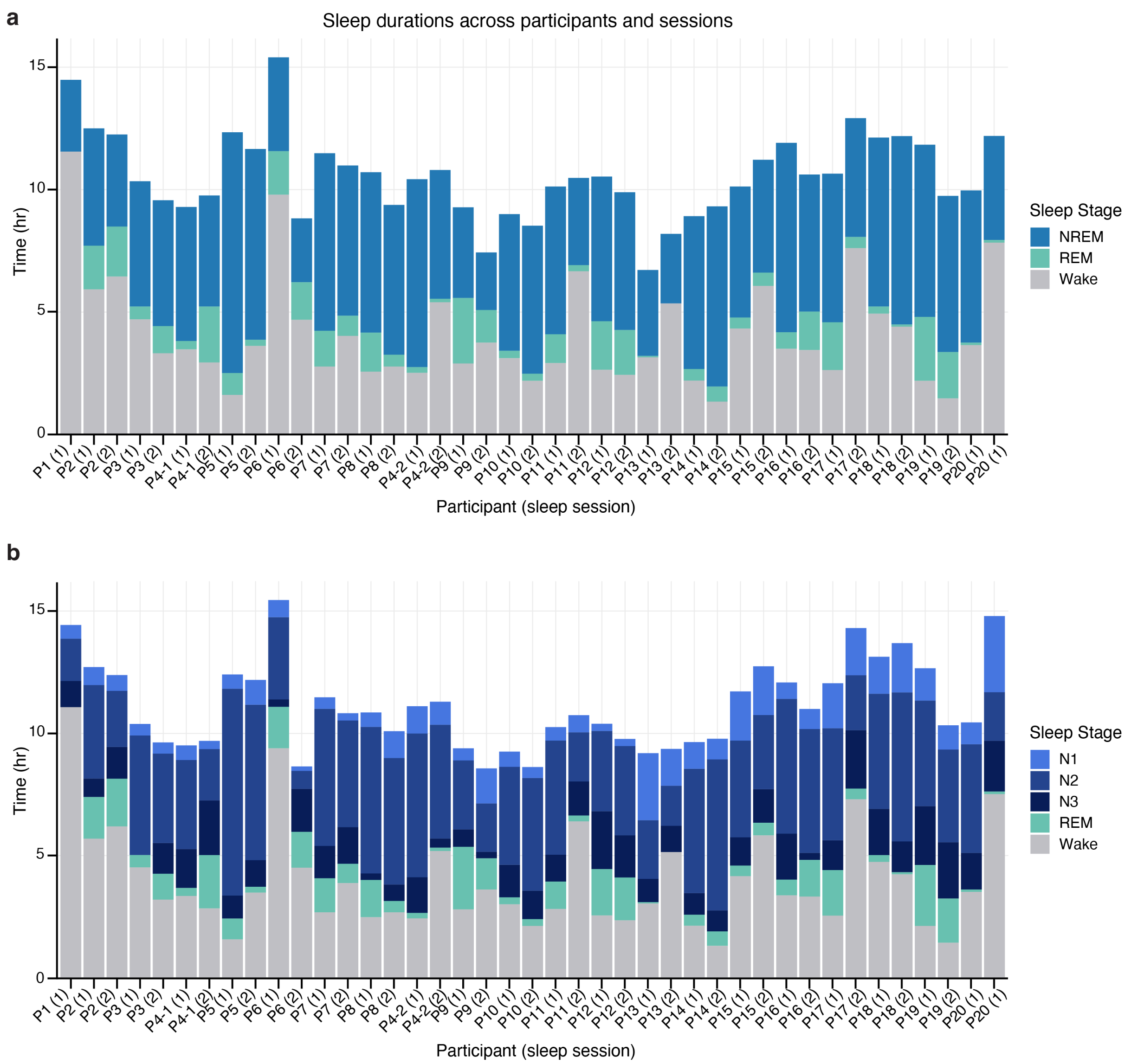


**Figure S1. Sleep stages and durations across participants. a.** Sleep stage and duration in hours for each participant and sleep session, stacked for total time in Wake, REM, and NREM (NREM = N2 and N3). **b.** Sleep stage and duration in hours for each participant and sleep session stacked for total time in Wake, REM, N1, N2, and N3. See Table S2 and S3 for individual participant and grand mean sleep durations by stage, respectively.

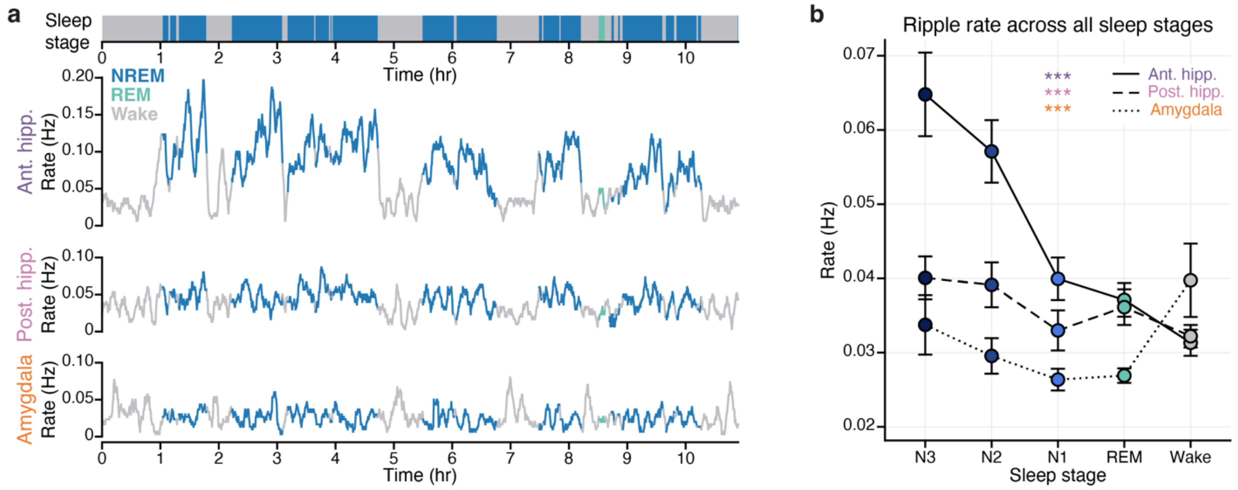


**Figure S2. Ripple rate across sleep stages—additional participant and stages. a.** Additional example of ripple rate across a night of sleep, for an electrode in anterior hippocampus (top), posterior hippocampus (middle), and amygdala (bottom), from one participant and hemisphere (P20, right hemisphere; same participant and sleep session as in Fig. 4a). Top bar: sleep stage over time (NREM: dark blue; REM: green; Wake: grey). **b.** Group mean ripple rate per sleep stage for all sleep stages (N1, N2, N3, REM, W), separately for electrodes in anterior hippocampus (solid), posterior hippocampus (dashed), and amygdala (dotted). Filled circles: group mean; error bars: ±SEM across group; significance of main effect of sleep stage: ***$p < 0.001$.

For Fig. S2b, the effect of all sleep stages on ripple rate was evaluated with a mixed-effects model, containing sleep stage (N3, N2, N1, REM, Wake) and region (anterior hippocampus, posterior hippocampus, amygdala) as fixed effects and electrodes nested within participants as random effects. Sleep stage, region, and the sleep stage × region interaction were all significant (sleep stage: $\chi^2(4) = 709.72$, $p < 0.001$; region: $\chi^2(2) = 210.41$, $p < 0.001$; sleep stage × region: $\chi^2(8) = 395.03$, $p < 0.001$). In anterior hippocampus, ripple rates were highest in N3 and lowest in wake, decreasing from N3 to N2 ($t(1698) = 7.12$, $p < 0.001$), N2 to N1 ($t(1698) = 10.58$, $p < 0.001$), and N1 to wake ($t(1698) = 4.81$, $p < 0.001$). N1 rates were not different from REM rates ($t(1699) = 2.11$, $p = 0.216$) and REM rates were not different from wake rates ($t(1699) = 2.66$, $p = 0.061$; $F(4, 1710.10) = 176.17$, $p < 0.001$). In posterior hippocampus, rates were highest in N3 and N2 and lowest in N1 and wake; ripple rates in N3 and N2 were both elevated relative to N1 (N3–N1: $t(1698) = 3.40$, $p = 0.006$; N2–N1: $t(1698) = 2.81$, $p = 0.040$) and wake (N3–wake: $t(1698) = 3.56$, $p = 0.004$; N2–wake: $t(1698) = 2.97$, $p = 0.025$). Rates in N3 and N2 did not differ from one another ($t(1698) = 0.61$, $p = 0.974$) and rates in N1, REM, and wake did not differ from one another (N1–REM: $t(1701) = 0.96$, $p = 0.874$; N1–wake: $t(1698) = 0.16$, $p = 0.9998$; REM–wake: $t(1701) = 1.12$, $p = 0.798$; $F(4, 1709.86) = 5.23$, $p < 0.001$). In amygdala, rates in wake and N3 were elevated relative to N2 (wake–N2: $t(1698) = 3.98$, $p = 0.001$; N3–N2: $t(1698) = 3.39$, $p = 0.007$), N1 (wake–N1: $t(1698) = 6.13$, $p < 0.001$; N3–N1: $t(1698) = 5.51$, $p < 0.001$), and REM (wake–REM: $t(1700) = 6.03$, $p < 0.001$; N3–REM: $t(1701) = 5.42$, $p < 0.001$), but not different from each other (N3–wake: $t(1698) = 0.55$, $p = 0.982$). N2, N1, and REM rates did not differ from one another (N2–N1: $t(1698) = 2.15$, $p = 0.200$; N2–REM: $t(1700) = 2.09$, $p = 0.225$; N1–REM: $t(1700) = 0.04$, $p = 1.000$; all Tukey-corrected; $F(4, 1710.09) = 16.88$, $p < 0.001$).

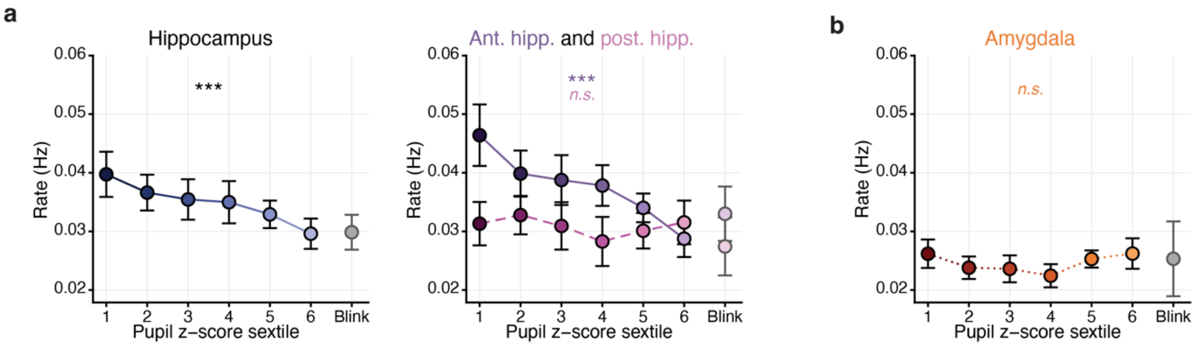


**Figure S3. Ripple rate across pupil states and blink window. a.** Group mean ripple rate across pupil states with the inclusion of a blink window, for all hippocampal electrodes (left) and separately for anterior hippocampus (solid) and posterior hippocampus (dashed) (right). The blink window represents the moment of blink occurrence plus 200ms after. **b.** Group mean ripple rate across pupil states with the blink window for all amygdala electrodes. Throughout, filled circles: group mean; error bars: ±SEM across group; significance of main effect of pupil sextile: ***$p < 0.001$.

After excluding ripples occurring during blinks (i.e., blink window ripples), the effect of pupil size on the remaining ripples was evaluated with a mixed-effects model predicting ripple rate as a function of pupil sextile (1–6) and region (anterior hippocampus, posterior hippocampus, amygdala), with participant and electrode as random effects. Pupil sextile, region, and the pupil sextile × region interaction were all significant predictors of ripple rate (pupil sextile: $\chi^2(1) = 19.87$, $p < 0.001$; region: $\chi^2(2) = 45.69$, $p < 0.001$; pupil sextile × region: $\chi^2(2) = 15.35$, $p < 0.001$). Anterior hippocampus ripple rate significantly tracked pupil sextile, decreasing as pupil size increased (slope = −0.00213 ± 0.00048 Hz, t(977) = −4.46, $p < 0.001$). Posterior hippocampus and amygdala showed no significant relationship to pupil sextile (posterior hippocampus: slope = −0.00018 ± 0.00056 Hz, t(977) = −0.32, $p = 0.748$; amygdala: slope = 0.000368 ± 0.000453 Hz, t(977) = 0.81, $p = 0.417$). The slope of the negative relationship between anterior hippocampus ripple rate and pupil sextile was significantly steeper than that of both posterior hippocampus (t(977) = −2.65, $p = 0.023$) and amygdala (t(977) = −3.80, $p = 0.001$; posterior hippocampus vs. amygdala: t(977) = −0.76, $p = 0.728$; all Tukey-corrected).

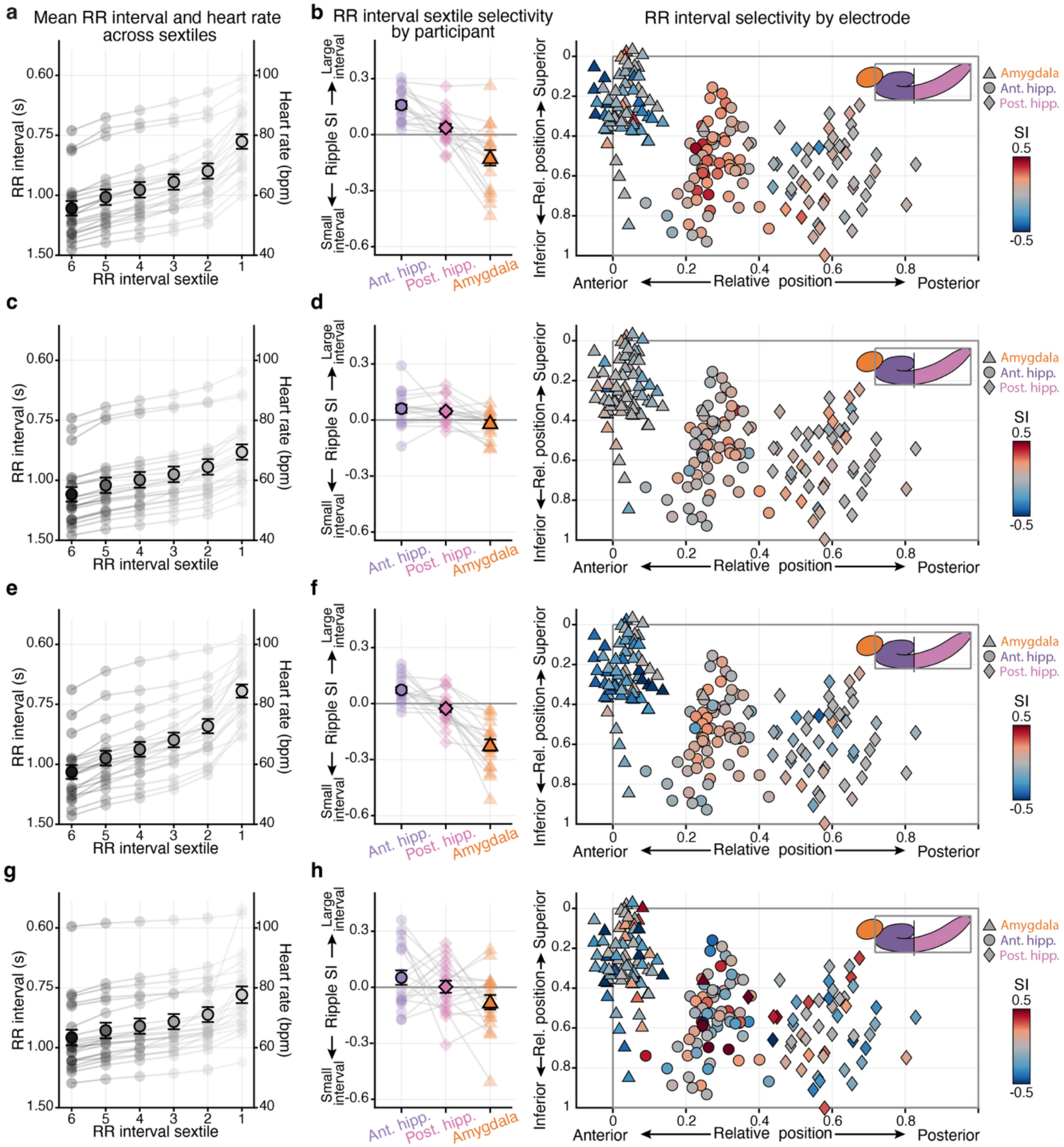


**Figure S4. Ripple rate selectivity to RR interval sextiles across sleep and wake. a.** Mean RR interval and heart rate per RR interval sextile during sleep. **b.** Ripple rate selectivity index (SI) to RR interval sextile during sleep. Positive values indicate selectivity to large RR interval, slow heart rate states (more ripples during large RR intervals than during small RR intervals) and negative values indicate selectivity to small RR interval, fast heart rate states. Left: SI by anatomical region for individual participants (transparent shapes) and group mean (solid shapes). Right: individual electrodes plotted by anatomical position relative to the most anterior and superior point of each participant's hippocampus, colored by SI (red: selectivity to large interval, slow heart rate; blue: selectivity to small interval, fast heart rate); shapes indicate anatomical region (anterior hippocampus: circle; posterior hippocampus: diamond; amygdala: triangle). **c–h ,** as in a, b for the NREM stages of sleep (c, d), wake stages of sleep (e, f), and wake fixation task (g, h), respectively. For **a, c, e, g** filled circles: group mean; error bars ±SEM across group; transparent dots and lines: individual participants.

The relationship between heart rate (RR interval sextile) and ripple rate across the continuous hippocampal long axis was evaluated in two ways. First, a selectivity index (SI) was calculated in a similar manner to that for NREM sleep and small pupil states (see main text). For each electrode, we quantified the degree to which ripple rates increased during low heart rate, large RR interval sextiles (low heart rate = average rate of sextiles 5 and 6) compared to high heart rate, small RR interval sextiles (high heart rate = average rate of sextiles 1 and 2): SI = (low heart rate – high heart rate) / (low heart rate + high heart rate). Positive and negative values reflect greater ripple rates during low heart rate and high heart rate states, respectively. Second, a mixed-effects model predicting ripple rate as a function of RR interval sextile and continuous anterior-to-posterior (AP) position was applied, where a significant interaction between RR interval sextile and AP position would indicate a change in the heart rate-ripple rate relationship across the long axis.

Across sleep, increases in ripple rate in low heart rate states were strongest in the most anterior hippocampal electrodes with SI values decreasing posteriorly (Fig. S4b). This effect was significant in both analysis of SI (SI slope (anterior to posterior) = −0.3102 ± 0.0566, t(106.34) = −5.478, $p < 0.001$) and in the full mixed-effects model (RR sextile × AP position = 0.0087 ± 0.0009, t(575) = 9.27, $p < 0.001$). In NREM only, increases in ripple rate in low heart rate states were strongest anteriorly, as measured by an interaction in the full mixed-effects model, although this result was not significant in analysis of SI (RR sextile × AP position = 0.0027 ± 0.0009, t(575) = 3.10, $p = 0.002$; SI slope = −0.0121 ± 0.034 t(99.655) = 0.353, $p = 0.725$; Fig. S4d). In the wake periods of the sleep recordings, increases in ripple rate in low heart rate states were again strongest in the most anterior hippocampal electrodes, measured both by a significant RR sextile × AP position interaction and analysis of SI values (RR sextile × AP position = 0.0039 ± 0.0006, t(575) = 6.132, $p < 0.001$; SI slope = −0.222 ± 0.0546, t(114.87) = −4.062, $p < 0.001$; Fig. S4f). Finally, the same trends of low heart rate states favoring ripple genesis especially in the most anterior hippocampal electrodes were observed in analysis of the wake fixation task, where a significant RR sextile × AP position interaction was found, although analysis of SI did not reach significance (RR sextile × AP position = 0.0041 ± 0.0018, t(575) = 2.275, $p = 0.023$; SI slope = −0.2395 ± 0.123 t(118.83) = −1.948, $p = 0.054$; Fig. S4h).

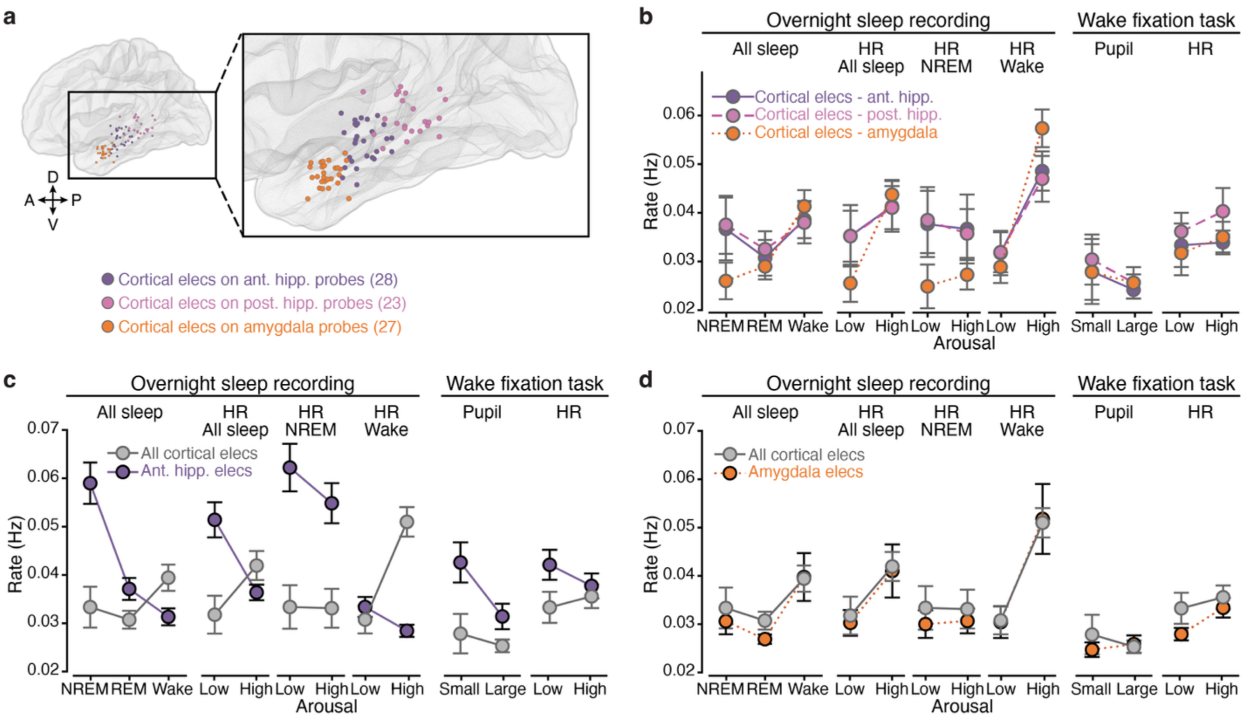


**Figure S5. Ripple-like burst activity in lateral temporal cortex across arousal states. a.** Intracranial electrode locations of lateral temporal electrodes for all participants normalized to MNI space, viewed from the lateral surface (hemispheres collapsed). Colors indicate the target of the probe that each electrode was on (purple = electrode on probe targeting anterior hippocampus, pink = electrode on probe targeting posterior hippocampus, orange = electrode on probe targeting amygdala). **b.** Group mean ripple rate across sleep stage, heart rate, and pupil arousal states for lateral temporal electrodes, separated by the region that their probe was targeting. **c.** Group mean ripple rate across sleep stage, heart rate, and pupil arousal states for all lateral temporal electrodes (grey) and for anterior hippocampus electrodes (purple). **d.** Group mean ripple rate across sleep stage, heart rate, and pupil arousal states for all lateral temporal electrodes (grey) and for amygdala electrodes (orange). For **b, c, d** heart rate states: low = mean of RR interval sextiles 5 and 6 (slow heart rate), high = mean of sextiles 1 and 2. Pupil states: small = mean of pupil sextiles 1 and 2, large = mean of sextiles 5 and 6; solid circles: group mean; error bars: ±SEM across group.

To evaluate whether lateral temporal cortex ripple-like events were modulated by arousal, and how their patterns related to that observed in the hippocampus and amygdala, mixed-effects models predicting ripple rate as a function of arousal (sleep stage: NREM, REM, wake; pupil sextile: 1–6; RR interval sextile: 1–6) and region (hippocampus, amygdala, cortex), with intercepts for electrodes nested within participants, were applied. Across sleep stages, pupil sextiles, and RR interval sextiles, results from the cortex differed from the hippocampus and aligned with the amygdala (Fig. S5c, d). During sleep, ripple-like event rate in cortex was significantly predicted by sleep stage, with rates highest during wake ($F(2, 1322.83) = 15.818$, $p < 0.001$). Pairwise comparisons revealed that the hippocampus differed significantly from the cortex across all sleep stages with the direction reversing between NREM and wake—the hippocampus had higher ripple rates than cortex during NREM (hippocampus – cortex: $t(489) = 8.99$, $p < 0.001$) and lower ripple rates during wake (cortex: $t(489) = -3.42$, $p = 0.002$); REM: cortex: $t(509) = 2.31$, $p = 0.055$; Fig. S5c). In contrast, the amygdala and cortex did not differ from one another at any sleep stage (all $t < 0.44$, $p > 0.90$; Fig. S5d). During the wake fixation task, ripple-like event rate in cortex was not significantly predicted by pupil sextile (slope (small to large pupil) = –0.0004 ± 0.0004 Hz, $t(1367) = -0.942$, $p = 0.346$). While the slope of this effect was not significantly different from that in the hippocampus (hippocampus – cortex = –0.0008 ± 0.0006 Hz, $t(1367) = -1.294$, $p = 0.399$), event rates in cortex were significantly lower than that in hippocampus at pupil sextile levels 1–5 and converged at level 6 (sextile 1: $t(699) = 4.110$, $p < 0.001$; sextile 2: $t(404) = 4.419$, $p < 0.001$; sextile 3: $t(271) = 4.460$, $p < 0.001$; sextile 4: $t(271) = 4.023$, $p < 0.001$; sextile 5: $t(404) = 3.236$, $p = 0.008$; sextile 6: $t(699) = 2.426$, $p = 0.090$; Šidák-corrected; Fig. S5c). The slope of cortex ripple-like event rate across pupil sextiles did not differ significantly from that of the amygdala either (amygdala – cortex = 0.0009 ± 0.0007 Hz, $t(1367) = 1.349$, $p = 0.369$), but, in contrast to the hippocampus, event rates in cortex were not different from amygdala at any pupil sextile level (all $|t| < 1.39$, all $p > 0.665$; Fig. S5d).

Examining the relationship between cortex ripple-like events and heart rate, across sleep, rates in cortex increased during fast heart rate states (small RR intervals; slope (slow to fast heart rate) = 0.0027 ± 0.0003 Hz, t(1318) = 10.082, $p < 0.001$). This pattern differed from that in the hippocampus, where event rates decreased during fast heart rate states (hippocampus – cortex = –0.0053 ± 0.0003 Hz, t(1318) = –15.427, $p < 0.001$; RR interval sextile 1: t(334) = 8.331, $p < 0.001$; sextile 2: t(278) = 6.311, $p < 0.001$; sextile 3: t(251) = 3.990, $p < 0.001$; sextile 4: t(251) = 1.508, $p = 0.575$; sextile 5: t(278) = –0.950, $p = 0.920$; sextile 6: t(334) = –3.215, $p = 0.009$; Fig. S5c). In contrast, event rates in the cortex and amygdala did not differ across heart rate levels (amygdala – cortex = –0.0002 ± 0.0004 Hz, t(1318) = –0.468, $p = 0.886$; across sextiles all |t| < 0.29, all $p > 0.999$; Šidák corrected; Fig. S5d). In NREM only, ripple-like events in cortex were not significantly predicted by RR interval sextile (slope = –0.00005± 0.0002 Hz, t(1318) = –0.342, $p = 0.733$), again differing significantly from the hippocampus where event rates decreased during fast heart rate states (hippocampus – cortex = –0.0016 ± 0.0002 Hz, t(1318) = –8.072, $p < 0.001$; RR interval sextile 1: t(261) = 6.694, $p < 0.001$; sextile 2: t(253) = 6.310, $p < 0.001$; sextile 3: t(249) = 5.897, $p < 0.001$; sextile 4: t(249) = 5.459, $p < 0.001$; sextile 5: t(253) = 5.001, $p < 0.001$; sextile 6: t(261) = 4.529, $p < 0.001$; Fig. S5c). Event rates in cortex and amygdala did not differ across heart rate levels within NREM (amygdala – cortex = 0.0002 ± 0.0002 Hz, t(1318) = 1.07, $p = 0.531$; across sextiles all |t| < 0.28, all $p > 0.999$; Šidák corrected; Fig. S5d). In wake only, ripple-like events in cortex dramatically increased during fast heart rate states (small RR interval sextiles; slope = 0.0052 ± 0.0003 Hz, t(1318) = 19.90, $p < 0.001$), differing significantly from the hippocampus (hippocampus – cortex = –0.0057 ± 0.0003 Hz, t(1318) = –16.89, $p < 0.001$; RR interval sextile 1: t(351) = 3.535, $p = 0.003$; sextile 2: t(283) = 0.856, $p = 0.950$; sextile 3: t(251) = –2.085, $p = 0.208$; sextile 4: t(251) = –5.051, $p < 0.001$; sextile 5: t(283) = –7.780, $p < 0.001$; sextile 6: t(351) = –10.089, $p < 0.001$; Fig. S5c) but converging with the amygdala across all heart rate levels (amygdala – cortex = 0.0001 ± 0.0004 Hz, t(1318) = 0.168, $p = 0.985$; across sextiles all |t| < 0.45, all $p > 0.997$; Šidák corrected; Fig. S5d). Finally, in the wake fixation task, mirroring that of the wake periods during sleep, ripple-like events in cortex increased during fast heart rate states (small RR interval sextiles; slope = 0.0010 ± 0.0004 Hz, t(1373) = 2.708, $p = 0.007$). Again, this effect differed significantly from hippocampus (hippocampus – cortex = –0.0014 ± 0.0005 Hz, t(1373) = –2.803, $p = 0.014$; RR interval sextile 1: t(585) = 2.966, $p = 0.019$; sextile 2: t(368) = 2.613, $p = 0.055$; sextile 3: t(271) = 2.011, $p = 0.243$; sextile 4: t(271) = 1.197, $p = 0.796$; sextile 5: t(368) = 0.354, $p = 1.000$; sextile 6: t(585) = –0.352, $p = 1.000$; Fig. S5c) but not from the amygdala (amygdala – cortex = 0.0007 ± 0.0006 Hz, t(1373) = 1.204, $p = 0.451$; across sextiles all |t| < 2.10, all $p > 0.201$; Šidák corrected; Fig. S5d).

**Table S1. Experimental data summary.** Summary of participant experimental data described across the manuscript, including electrode counts by region, nights of sleep, blocks of the wake fixation task, and number of eyes-open (EO) trials. *Participant #4 underwent two separate implants and recordings.

| Participant | Hippocampal electrodes | Ant. hipp. electrodes | Post. hipp. electrodes | Amygdala electrodes | Sleep nights | Fixation task blocks | EO trials |
|---|---|---|---|---|---|---|---|
| 1 | 5 | 2 | 3 | 3 | 1 | 4 | 37 |
| 2 | 6 | 4 | 2 | 6 | 2 | 6 | 88 |
| 3 | 7 | 5 | 2 | 6 | 2 | 4 | 48 |
| 4 - 1* | 2 | 0 | 2 | 3 | 2 | 1 | 14 |
| 4 - 2* | 4 | 3 | 1 | 4 | 2 | 4 | 52 |
| 5 | 4 | 4 | 0 | 0 | 2 | 4 | 48 |
| 6 | 5 | 3 | 2 | 3 | 2 | 4 | 44 |
| 7 | 8 | 6 | 2 | 5 | 2 | 5 | 49 |
| 8 | 4 | 2 | 2 | 3 | 2 | 4 | 39 |
| 9 | 10 | 6 | 4 | 0 | 2 | 6 | 69 |
| 10 | 3 | 1 | 2 | 4 | 2 | 3 | 29 |
| 11 | 9 | 4 | 5 | 3 | 2 | 9 | 122 |
| 12 | 5 | 3 | 2 | 3 | 2 | 9 | 94 |
| 13 | 4 | 2 | 2 | 2 | 2 | 4 | 40 |
| 14 | 5 | 2 | 3 | 3 | 2 | 8 | 80 |
| 15 | 10 | 8 | 2 | 6 | 2 | 3 | 23 |
| 16 | 2 | 0 | 2 | 5 | 2 | 6 | 80 |
| 17 | 4 | 3 | 1 | 2 | 2 | 4 | 45 |
| 18 | 9 | 4 | 5 | 7 | 2 | 4 | 44 |
| 19 | 3 | 1 | 2 | 6 | 2 | 5 | 49 |
| 20 | 10 | 6 | 4 | 3 | 2 | 4 | 39 |

| Total participants | Total hipp. electrodes | Total ant. hipp. electrodes | Total post. hipp. electrodes | Total amygdala electrodes | Total sleep nights | Median fixation task blocks (IQR) | Median EO trials (IQR) |
|---|---|---|---|---|---|---|---|
| 20 | 119 | 69 | 50 | 77 | 41 | 4 (4–6) | 48 (39–69) |

**Table S2. Sleep information for individual participants.** Time in minutes in each sleep stage, for each sleep session. Total sleep reflects time across REM, N1, N2, and N3. Total NREM is time in N2 and N3. "EKG recorded" indicates whether EKG was recorded and analyzed from each sleep session. *Participant #4 underwent two separate implants and recording periods.

| Participant | Session | Wake | REM | N1 | N2 | N3 | Total sleep | Total NREM | EKG recorded |
|---|---|---|---|---|---|---|---|---|---|
| 1 | 1 | 693 | 0 | 35 | 108.5 | 67.5 | 211 | 176 | yes |
| 2 | 1 | 355.5 | 107 | 46 | 240 | 47.5 | 440.5 | 287.5 | yes |
| 2 | 2 | 387 | 122.5 | 40.5 | 144 | 81.5 | 388.5 | 225.5 | yes |
| 3 | 1 | 282 | 31.5 | 29.5 | 307 | 0 | 368 | 307 | yes |
| 3 | 2 | 199 | 66.5 | 28.5 | 229.5 | 79 | 403.5 | 308.5 | yes |
| 4-1* | 1 | 208.5 | 20.5 | 37.5 | 228.5 | 100 | 386.5 | 328.5 | yes |
| 4-1* | 2 | 176.5 | 137 | 21 | 131.5 | 140.5 | 430 | 272 | yes |
| 4-2* | 1 | 151 | 14 | 70 | 369 | 91.5 | 544.5 | 460.5 | no |
| 4-2* | 2 | 324 | 8.5 | 59 | 292 | 23.5 | 383 | 315.5 | no |
| 5 | 1 | 97 | 53.5 | 36.5 | 531 | 59 | 680 | 590 | yes |
| 5 | 2 | 217 | 15 | 63.5 | 399 | 68.5 | 546 | 467.5 | yes |
| 6 | 1 | 587.5 | 106.5 | 44 | 211 | 19 | 380.5 | 230 | yes |
| 6 | 2 | 281 | 92 | 11.5 | 46 | 110.5 | 260 | 156.5 | yes |
| 7 | 1 | 166.5 | 87.5 | 29.5 | 352 | 83 | 552 | 435 | yes |
| 7 | 2 | 241.5 | 49.5 | 18.5 | 274 | 94 | 436 | 368 | yes |
| 8 | 1 | 154 | 95.5 | 37 | 376 | 17 | 525.5 | 393 | yes |
| 8 | 2 | 166.5 | 29 | 69 | 324.5 | 42.5 | 465 | 367 | yes |
| 9 | 1 | 174 | 160.5 | 31 | 177 | 45 | 413.5 | 222 | yes |
| 9 | 2 | 225 | 80 | 90 | 124 | 17 | 311 | 141 | yes |
| 10 | 1 | 187 | 18 | 39 | 251.5 | 83.5 | 392 | 335 | yes |
| 10 | 2 | 131.5 | 17.5 | 28 | 289.5 | 73 | 408 | 362.5 | yes |
| 11 | 1 | 175 | 70.5 | 34.5 | 293 | 69 | 467 | 362 | yes |
| 11 | 2 | 400 | 15 | 44 | 126 | 87.5 | 272.5 | 213.5 | yes |
| 12 | 1 | 158.5 | 119 | 18.5 | 206 | 148.5 | 492 | 354.5 | yes |
| 12 | 2 | 146 | 110 | 18 | 229.5 | 108 | 465.5 | 337.5 | yes |
| 13 | 1 | 189 | 3.5 | 172 | 150.5 | 60 | 386 | 210.5 | yes |
| 13 | 2 | 321 | 0 | 94.5 | 103 | 67.5 | 265 | 170.5 | yes |
| 14 | 1 | 132 | 28.5 | 68.5 | 319 | 55.5 | 471.5 | 374.5 | yes |
| 14 | 2 | 80.5 | 37 | 53 | 387.5 | 54 | 531.5 | 441.5 | yes |

| | | | | | | | | | |
|---|---|---|---|---|---|---|---|---|---|
| 15 | 1 | 259.5 | 27 | 126 | 248.5 | 72.5 | 474 | 321 | yes |
| 15 | 2 | 364 | 32.5 | 125 | 190.5 | 86 | 434 | 276.5 | yes |
| 16 | 1 | 210 | 40.5 | 42 | 346 | 118 | 546.5 | 464 | no |
| 16 | 2 | 207 | 94 | 51.5 | 318 | 18 | 481.5 | 336 | yes |
| 17 | 1 | 158 | 117 | 115.5 | 287.5 | 76.5 | 596.5 | 364 | yes |
| 17 | 2 | 456.5 | 27.5 | 121 | 141 | 150 | 439.5 | 291 | yes |
| 18 | 1 | 296 | 17.5 | 95 | 295.5 | 118.5 | 526.5 | 414 | yes |
| 18 | 2 | 264 | 5.5 | 126.5 | 382.5 | 79 | 593.5 | 461.5 | yes |
| 19 | 1 | 131.5 | 156.5 | 83 | 271 | 151 | 661.5 | 422 | yes |
| 19 | 2 | 88.5 | 113.5 | 62 | 238 | 144.5 | 558 | 382.5 | yes |
| 20 | 1 | 219 | 6 | 56 | 279 | 94 | 435 | 373 | yes |
| 20 | 2 | 470 | 6.5 | 195.5 | 124.5 | 130.5 | 457 | 255 | yes |

**Table S3. Sleep duration summary across participants.** Summary statistics for time (minutes) spent in different sleep stages across all participants and nights. NREM is time in N2 and N3. Total sleep is time in N1, N2, N3, and REM.

| Sleep stage | Mean | Standard deviation | Minimum | Maximum |
|---|---|---|---|---|
| Wake | 247.10 | 132.00 | 80.50 | 693 |
| N1 | 62.60 | 42.78 | 11.50 | 195.50 |
| N2 | 252.24 | 100.87 | 46 | 531 |
| N3 | 78.82 | 39.44 | 0 | 151 |
| REM | 57.06 | 47.89 | 0 | 160.50 |
| NREM | 331.06 | 98.35 | 141 | 590 |
| Total sleep | 450.72 | 103.45 | 211 | 680 |

**Table S4. Participant demographics.** Information on hemisphere of implant, as well as participant handedness, sex, age, and race.

| Participant | Hemisphere | Handedness | Sex | Age at Implant | Race |
|---|---|---|---|---|---|
| 1 | L | R | F | 59 | W |
| 2 | L, R | R | M | 30 | BA |
| 3 | L, R | R | M | 57 | W |
| 4 | L | L | M | 26 | W |
| 5 | L, R | R | F | 28 | BA |
| 6 | R | R | F | 28 | W |
| 7 | L, R | R | F | 29 | BA |
| 8 | L | R | M | 37 | BA |
| 9 | L, R | R | M | 46 | ME |
| 10 | L, R | L | M | 46 | HL |
| 11 | L, R | R | F | 61 | W |
| 12 | R | R | F | 34 | Unknown |
| 13 | R | R | M | 64 | BA |
| 14 | L | R | F | 46 | W |
| 15 | L, R | R | F | 21 | W |
| 16 | L, R | R | M | 27 | W |
| 17 | L | L | F | 44 | W |
| 18 | L, R | R | F | 50 | BA |
| 19 | L, R | R | M | 29 | AI |
| 20 | L, R | R | M | 33 | BA |

W: White; BA: Black or African American; AI: American Indian; HL: Hispanic or Latinx; ME: Mixed ethnicity
